# A full computational model of cell motility: Early spreading, cell migration and competing taxis

**DOI:** 10.1101/2022.09.28.509519

**Authors:** Joel Betorz, Gaurav R. Bokil, Shreyas M. Deshpande, Shardool Kulkarnic, Daniel Rolando, Chiara Venturini, Pablo Saez

**Affiliations:** ETS de Ingenieros de Caminos, Canales y Puertos; Laboratori de Cacùl Numèric (LaCàN); Facultat de Matemàtiques i Estadìstica; IMTech (Institute of Mathematics), Universitat Politècnica de Catalunya-BarcelonaTech., 08034 Barcelona, Spain

**Keywords:** Cell motility, mechanobiology, durotaxis, chemotaxis, tissue engineering

## Abstract

Cell motility represents one of the most fundamental function in mechanobiology. Cell motility is directly implicated in development, cancer or tissue regeneration, but it also plays a key role in the future of tissue and biomedical engineering. Here, we derived a computational model of cell motility that incorporates the most important mechanisms toward cell motility: cell protrusion, polarization and retrograde flow. We first validate our model to explain two important types of cell migration, i.e. confined and ameboid cell migration, as well as all phases of the latter cell migration type, i.e. symmetric cell spreading, cell polarization and latter migration. Then, we use our model to investigate durotaxis and chemotaxis. The model predicts that chemotaxis alone induces larger migration velocities than durotaxis and that durotaxis is activated in soft matrices but not in stiff ones. More importantly, we analyze the competition between chemical and mechanical signals. We show that chemotaxis rules over durotaxis in most situations although durotaxis diminishes chemotaxis. Moreover, we show that inhibiting the effect of GTPases in actin polymerization at the cell front may allow durotaxis to take control over chemotaxis in soft substrates. Understanding how the main forces in cell motility cooperate, and how a precise manipulation of external cues may control directed cell migration is not only key for a fundamental comprehension of cell biology but also to engineer better biomimetic tissues. To this end, we provide a freely-available platform to predict all phases and modes of cell motility analyzed in this work.

## 1 Introduction

Cell motility is a mechanical function central to life [1]. During development, cells migrate individually and collectively to, ultimately, determine the underlying mechanical structure and function of biological tissues and organs [2, 3]. Cell migration is also tightly implicated in diseases. For example, cell motility is responsible for tumor cells to invade healthy tissues and metastasize [4, 5]. Motile cells also orchestrate the regeneration or wounding of tissues, i.e. how tissues heal if they are damaged [1, 6]. Not only it is central to physiology and disease, cell motility is key for engineering novel biomimetic tissue designs [7, 8].

Understanding cell motility requires tackling several physical mechanisms that determine its different phases, from the early cell spreading [9] to the later polarization [10, 11] and the subsequent cell migration [1]. Cell motility is enabled by internal propelling machinery that consists of two main mechanisms. One is a continuous polymerization of actin filaments at the leading edge of the cell that protrudes the cell membrane forward [12, 13]. The other force arises from the pulling that myosin motors exert on the actin network, which generate an active retrograde flow of actin [14, 15]. Resisting the retrograde flow, a large number of cell adhesion molecules (CAMs) form attachment between cells and their surroundings [16]. CAMs, including integrins, talin, or vinculin, among others, link the retrograde flow with the extracellular space through a dynamic process of complex adhesion growth, and disassembly [16, 17]. Without adhesion, cells would not move because those intracellular forces could not be transmitted outside of the cell. Together, the retrograde flow, the actin protrusion against the cell membrane and the cell adhesions are the most important mechanical functions toward cell motility.

### 1.1 The acto-myosin network and its role in cell motility

Actin is unarguably a fundamental element in cell function, and it is specifically key in cell motility [18, 15, 19]. The actin network is a dynamic system that continuously assembles and disassembles to form unique cellular structures. At the very front of the cell, a network of actin filaments, forms and protrudes the cell membrane forward [20, 21, 19]. In the cell cortex and in the lamella, another dynamic and contractile network of actin filaments determines the fate of cell motility [21, 19].

The basic features of the turnover cycle of actin at different spatial and temporal scales of the cell is now fairly well understood [15, 22]. Actin monomers nucleate, promoted by Formin, Profilin and Cofilin [23, 24], and polymerize rapidly after a period of slow nucleation [19, 22]. A key actor in this process is the nucleation promoting factor Arp2/3. Arp2/3 activates and promotes the formation of new branches of actin filament along the barbed end of the actin filament [25]. Some of these actin polymerization promoters are activated by a number of GTPases (see Section 1.3 for details). Actin filaments, or F-Actin, are identified as a polar structure that acquires a fast-growing end, the barbed end, and a slow dissociating end, the pointed end. Barbed ends terminate its formation when capping proteins associate in a process referred as capping [26] inhibiting further growth. F-Actin dissociates, mediated by ADP and cofilin, at the pointed-end producing ADP-actin dissociations. Then, profilin promotes the ATP replacement of ADP, providing a pool of new actin monomers ready to bind the barbed ends [15, 22].

In order to regulate the retrograde flow of the F-actin network required for cell motility [27, 28, 29], myosin motors pull on the individual actin filaments. The motor domain of the myosin protein, a superfamily of ATP-dependent motor proteins, binds to actin filaments and generates force via ATP hydrolysis to move along the filament towards the barbed end [30]. These forces generate an asymmetric retrograde flow that polarizes the actomyosin network by convective forces. This polarization of the actomyosin network is partially responsible for cell motility [31]. The force generated by the myosin motors is also implicated in the cell adhesion dynamics through a number of mechanosensing mechanisms in the CAMs [32]. Therefore, the actomyosin network is not only responsible for the active movement of the F-actin network, and therefore for the generation of an active motile force, but it also controls cell adhesion dynamics.

### 1.2 Early symmetric cell spreading

One of the very first motile phases is cell spreading [9], an expanding, flattening process in which cells polarize and acquire motile features. Importantly, cell spreading is highly reproducible and, therefore, a perfect system to analyze the basic mechanisms of cell motility in space and time in a highly quantitative and predictable manner. Yet, how biochemical and biophysical forces coordinate cell spreading is not clear. What is now widely accepted is that cell spreading, as a form of cell motility, is coordinated by the active and contractile actomyosin network [33, 34], the actin polymerization at the cell front [20, 21, 19], cell adhesion mechanisms [16, 35] and membrane tension [36, 37, 38].

Prior to the cell polarization, a symmetric cell spreading occurs within 10-20 minutes after the cells have been seeded on the substrate [9] and consists of three main phases: 1) cells initially attaching to the ECM (phase P0) by integrin-based adhesions, with constant contact area; 2) then they rapidly spread (phase P1) increasing the contact area considerably; and, finally, 3) cell spreading slows down until they reach a steady state. All the phases are distinguished by specific signatures of acto-myosin flows, actin protrusions and adhesion mechanisms.

In phase P0, the cell attaches to the ECM due to the integrin clustering at the cell periphery [39, 40, 41] which depends on the mechanosensing of CMAs, including integrins [35] and talin [42, 43]. Once the nascent adhesions have taken place, the activation of the motile forces initiates, and a fast spreading phase, P1, starts. The fast spreading at P1 is driven by the polymerization of actin against the cell membrane. Moreover, after the transient nascent adhesions form in P0, they either disintegrate or grow into mature focal adhesions [44, 45, 46] by the end of P1 [47]. During P1, the cell membrane is capable of increasing its area via endocytosis and by progressively depleting the membrane reservoirs [48]. Once all membrane reservoirs are consumed, the cell membrane tension starts to increase by the end of P1, when the rapid spreading in P1 decelerates. Although not fully understood, it is believed that biochemical signals, prompted by physical forces imposed by the plasma membrane [39, 48, 37, 49], induce changes in actin polymerization, exocytosis [48, 49] and myosin activity.

### 1.3 Signaling and cell polarization

Before the cells crawls by itself, an external cue is required for the cells to polarize and establish a directed migration. To achieve a polarized state, cells need external stimuli in the form of, e.g., shallow chemical gradients, localized signals on the cell membrane or of mechanical stimuli. Indeed, triggered by some of those stimuli, different signaling cues polarizes at some point during the symmetric cell spreading phase. Then, they regulate the downstream polarization of the actomyosin network to, eventually, stablish a directed cell migration [10, 11]. There are many proteins involved in this process. Specially relevant are the members of the Rho GTPases family [50, 10, 51, 52]. Upstream, a positive-feedback loop between phosphati-dylinositol 3,4,5-trisphosphate (PIP3) and GTPase exchange factors (GEFs) [53] seems to be responsible for Rho GTPases family alterations. Rho GTPases includes the well characterized, and specifically involved in cell polarization, Cdc42 [54, 55, 56, 57], Rac [53] and RhoA [58]. Actin polymerisation against the cell membrane is promoted in regions of high Cdc42 and Rac1 concentration [53, 51, 59, 60, 10], which acts on downstream regulators of actin branching, including phosphoinositide 3-kinase (PI3k), WAVE proteins and Arp2/3 [61, 62, 63, 64, 33]. On the other hand, the downstream alterations in RhoA regulates membrane exocytosis and myosin activation, which eventually increase the myosin activity.

Moreover, Rac1 and RhoA are mutually inhibitory [65, 66]. Therefore, motile cells respond to the GTPases polarization by establishing two well-defined regions, the front and the back. The front is characterized by high levels of Rac1 and Cdc42, and higher rates of F-actin protrusions, while the rear is less active in terms of actin polymerisation and has more active myosin motors. Cells migrate toward the direction in which the front forms, where the actin polymerization protrudes the cell membrane, retracting the cell rear and, eventually, translocating the cell body in that same direction.

### 1.4 Modes and taxis in cell migration

Once cells break the spreading symmetry, they establish a rear-front profile and start to migrate in a specific direction. Cells migrate following distinct physical modes of locomotion which can be generally categorized as ameboid and mesenchymal cell migration [67, 68]. Mesenchymal cell migration is driven by the competition of actin protrusions against the cell membrane and the inward retrograde flow, expressed in the lamella and lamellipodia [21], which allows the cell front to extend forward and the cell rear to retract. Mesenchimal cell migration is also characterized by strong adhesions with the ECM. On the other hand, ameboid cell migration is characterized by the lack of a protruding actin network. The forward movement is performed by pseudopods and blebs, that squeeze through empty spaces of the matrix, and it is accompanied by the contractility of a highly polarized actomyosin cortex. Moreover, ameboid migration expresses low friction with the environment because strong, integrin-based, adhesion complexes are weakly expressed. There are other modes of cell migration, such as lobopodia [67], an intermediate mode between the amoeboid and mesenchymal modes, or confined migration, where a low-adhesion between the moving cell and its surroundings appears through simple contact pressure [69, 70] and cells migrate slowly due to transcellular water fluxes mediated by the chemical balance of the cell [71].

Besides random, inherent exogenous cues, cells can respond to externally imposed signals. Indeed, the response of cells to external stimuli during migration has been studied in cell biology for decades. Chemical and mechanical cues for cell motility, usually referred to as chemotaxis [72, 73] and durotaxis [74, 75], respectively, are probably the most studied forms of cellular taxis. During chemotaxis, the accumulation of chemical concentrations close to the cell membrane promotes downstream signaling mechanisms. Cells sense chemical signals through a number of membrane receptors [72] that establish an specific polarization direction of intracellular signals, e.g. GTPases, and, eventually, triggers the downstream polarization of the motile machinery of the cell [76, 10]. Durotaxis, or the use of stiffness gradients, has also been proved on different cells and tissues as well as in single and collective cell migration [3, 74, 75]. Overall, chemical and mechanical stimuli impose macroscopic signals that modify the motile mechanisms of cell migration.

### 1.5 Goal and structure of the manuscript

Mathematical and computational models have been extensively used (see, e.g., [77, 78, 79, 80] for a review) to unravel open questions in cell motility. The mathematical foundations of actin treadmilling have been well established [81, 82, 83, 84] and more and more detailed models appeared [85, 86, 83]. Some models coupled the actin polymerization with the mechanics of the membrane protrusion [87, 20, 88]. Other types of models described the retrograde flow and established the basic ingredients to describe the mechanics of the actomyosin network [89, 90, 34]. More recent computational models described the dynamics of the actin polymerization at the cell front while also accounting for the retrograde flow and the changes in the densities of the different constituents of the actomyosin network [91]. Some of the most detailed theoretical models today (see, among others, [82, 83]) were solved either analytically or in simple 1D numerical schemes. Others were solved by standard finite element approaches, phase-field models [92] and immersed boundary methods [93], which allow to describe complex shape changes in 2D and 3D. Most of the most complete models have been used to understand specific modes of migration, to reproduce specific stages of the cell migration process or certain taxis.

However, a single mechanistic model that describes all main modes and phases of cell motility, and that uniquely describes the underlying physical forces, has not been explicitly reported. Moreover, although different migration models have tackled durotaxis [74, 94] and chemotaxis [95, 96, 97, 98, 99], how chemical and mechanical signals compete to guide cell migration has not been investigated, and only isolated data have been reported [100]. This is a key aspect in cell migration, because stiffness gradients and chemical signals coexist in in-vivo systems. In this work, we integrate theoretical models of actin protrusion and the actomyosin network to develop a fully coupled finite element model of cell motility. The goal of this work is to first develop a single and complete model of cell motility that it incorporates all main mechanisms of cell motility and that reproduces every single phase and mode of cell migration. We use the model to reproduce previous works on chemotaxis and durotaxis and provide a rational description of these taxis. Finally, we analyze how chemical and mechanical cues compete during cell migration.

The paper is organized as follows. First, we detail the basic model ingredients, describing the transport mechanisms and the mechanics of the actomyosin network. We also summarize the numerical implementation by using the finite element method, where the temporal and spatial discretization are discussed. Then, we validate our model to explain the main phases of cell migration, i.e. symmetric cell spreading, cell polarization and later migration. We focus on two important types of cell migration, i.e. mesenchymal and ameboid cell migration. Then, we use the model to investigate durotaxis and chemotaxis and provide a physical rationale of the underlying forces and the dynamics of both processes. More importantly, we investigate the competing forces that control a specific directional response when chemotaxis and durotaxis appear simultaneously, which has been studied before neither experimentally nor mathematically. Finally, we discuss our results and conclude.

## 2 A full model of cell motility

We consider a 1D domain Ω (see Fig. 1) with moving coordinates *x*(*t*) *∈* [*l*_*r*_(*t*), *l*_*f*_ (*t*)] that integrates main ingredients for cell motility [79]. The subindexes *r* and *f* indicate the front and rear of the cell. *l*_*r*_(*t*) and *l*_*f*_ (*t*) represent the rear and front boundary of the cell and, therefore, the length of the cell is determined as *L*(*t*) = *l*_*f*_ (*t*) *− l*_*r*_(*t*). Both boundaries move with velocity *l*?_*r*_(*t*) and *l*?_*f*_ (*t*) and the migration velocity of the cell is *V*_*cell*_ = (*l*?_*r*_(*t*) + *l*?_*f*_ (*t*))*/*2. To model the mechanisms that control cell motility, we consider the transport of myosin motors and an actin network to describe how they localize in the cell body. We model the mechanics of the acto-myosin network as a viscous F-actin network that shows contractility due to the pulling forces exerted by myosin motors [90, 101, 102]. Then, we include a model of cell membrane protrusion driven by actin polymerization at both sides of the cell [103]. This actin polymerization depends simultaneously on the cell membrane tension [37, 49] and on the signaling of GTPases, which also regulate myosin activity. We also include the polarization of the GTPases [104].

**Figure 1:**
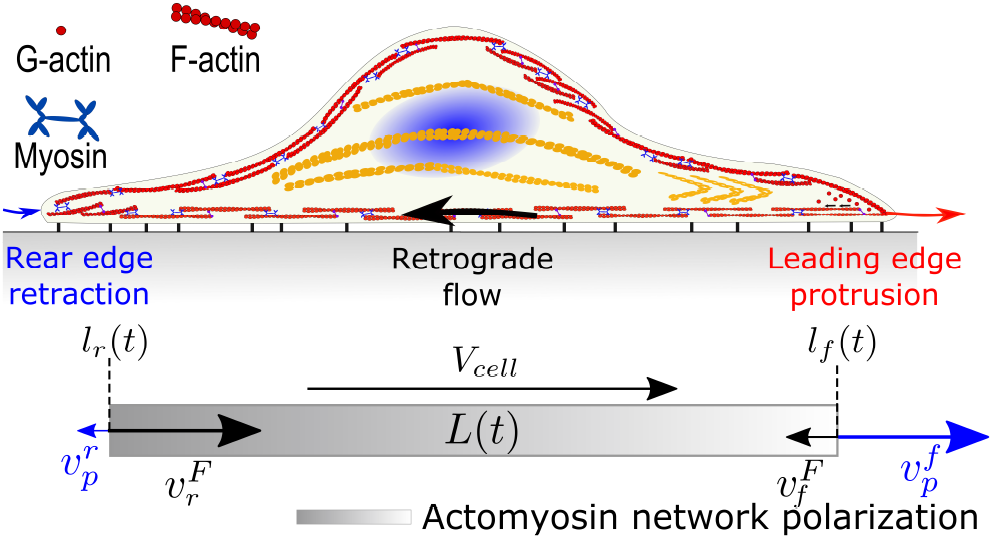
Sketch of the migration model and the forces acting to move the cell. A network of actin filaments pushes the cell membrane with polymerization velocity, *v*_*p*_. A second actin network fuses with myosin motors to create a contractile actomyosin network. As a results, a retrograde flow of the actomyosin network flows with velocity *v*^*F*^. If *v*_*p*_ becomes larger at the front than at the rear or the actomyosin network polarizes, then the cell undergoes a directed cell migration and controls the cell migration velocity, *V*_*cell*_. The difference of these two velocities at the cell fronts determines the cell length, *L*(*t*).

### 2.1 Transport of the F-actin network

To model the distribution in time and space of the actin density, we consider two advection-diffusion reaction equations that describe the F-actin network density, *ρ*^*F*^ (*x, t*), as well as the monomeric form of actin, G-actin density, *ρ*^*G*^(*x, t*). Therefore, we model the effective transport of these two actin forms as

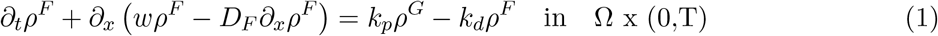

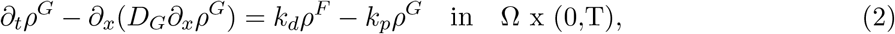

where the source term involves the polymerization and depolymerization rates, *k*_*p*_ and *k*_*d*_, respectively. The G-actin only diffuses because diffusion dominates the convection effects of the intracellular fluid where monomers drift. *D*_*F*_ and *D*_*G*_ are the diffusive parameters of the F-actin and G-actin, respectively. We write the transport equations in the cell frame and, accordingly, we define the velocity of the F-actin network in the cell frame as *w* = *v*^*F*^ *− V*_*cell*_, where *v*^*F*^ is the velocity of the actin network in the lab frame of reference (see Section 2.4). We impose zero fluxes on the boundary of the domain to reflect that no actin can enter or leave the cell domain.

A two species model of the actin turnover is a simplification of the actual actin dynamics [91, 15] (see also Section 1.1). Although more complete actin turnover models have been proposed [91, 105], the two-species model is a good approximation to model the effective transport of the F-actin and G-actin distribution [91]. Many other models have focused on a single transient advection-diffusion problem with first order kinetics for the F-actin phase, as in Eq. 1, that also showed good prediction of experimental findings [106, 91, 102]. This approach assumes that the concentration of G-actin is constant in space due to a fast diffusion process [106, 91] but changes in time as 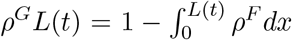. We analyze these two considerations in the modeling of cell motility in the following sections.

### 2.2 Transport of myosin motors

To describe the distribution in space and time of the myosin motors density, we also consider a two species model of bound, *ρ*^*M*^, and unbound, *ρ*^*m*^, myosin motors to the F-actin network [90, 102]. We model a system of two transient convection-diffusion reaction equations as

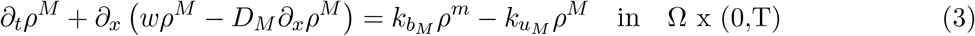

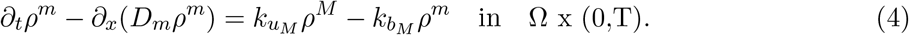

*D*_*M*_ and *D*_*m*_ are the diffusion parameters of the bound and unbound species, respectively. The transport of the unbound motors is governed by a diffusive term, because diffusion dominates convective effects. The myosin species also have to obey zero flux boundary conditions to describe that no myosin motors can enter or leave the cell.

Myosin motors unbind from the F-actin network with a constant rate 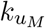, diffuse in the cytoplasm and bind again to the F-actin with a rate *k*_*bM*_ [107]. If one assumes that the binding-unbinding process is fast, i.e., *k*_*uM*_ */k*_*bM*_ *<<* 1 [102], we can model the effective transport of bound myosin motors, *ρ*^*M*^, through one single transient advection-diffusion problem [90, 108, 102] that advects the bounded myosin motors with velocity *w* as

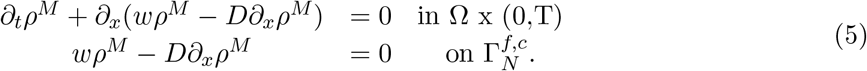

where 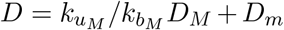 is the effective diffusion parameter [108, 102] and 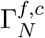 are Neumann boundary condition at the front and rear of the cell. Both forms of describing the myosin dynamics for cell migration are also discussed in following sections.

### 2.3 Polarization of signaling cues

During spreading (see Section 1.2), cells receive external cues that they use to polarize intracellular signals and to, directly or indirectly, mediate in the motile forces of the cells. Eventually, cells orientate themselves towards a target direction. This process is commonly refered to as polarization [10, 11, 53].

To achieve a polarized state, cells are exposed to exogenous stimuli in the form of, e.g., a shallow or a highly localized chemical or mechanical signals. These signals are sensed and transmitted by cell membrane receptors that later transmit the information intracellularly and respond by reorganizing the cell through two well-defined regions, front and back [10]. This polarization results in an asymmetric expression of the motile forces of the cell.

There are many proteins involved in cell polarization, including PI3K, PTEN and members of Rho GTPases family [50, 10, 51, 52] (see details in Section 1.3), which are ubiquitous in many cell types. Every Rho GTPase cycles between the plasma membrane (active GTP-bound form) and the cytosol (inactive GDP bound form). The active forms of Rho GTPases are responsible for activating downstream changes in the motile machinery of the cell.

Here we follow a well established model that simplifies the problem to one single GTPase in its active and inactive form, without further consideration of specific members of this family (see [104] for details). The so called wave-pinning model, which describes essential features of cell polarization, relies on two reaction diffusion equations with bistable kinetics and a positive feedback from the activation form onto its own production. The equations for the active GTPase density *ρ*^*R*^ and inactive form *ρ*^*S*^ are

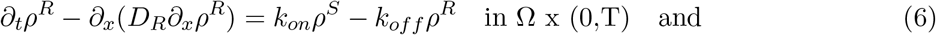

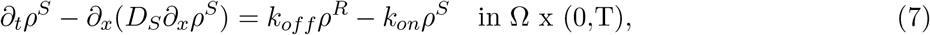

where *k*_*on*_ and *k*_*off*_ are the activation and inactivation rates, respectively. The diffusion rate of the active form is known to be significantly smaller than its inactive cytosolic counterpart, i.e. *D*_*R*_ ≪ *D*_*S*_. Here we follow a simple positive feedback from the activated form to its own production and also a constant inactivation rate [104],

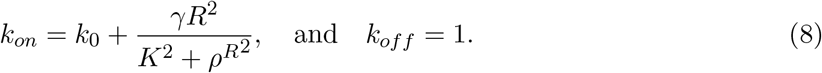

*k*_0_ = 1*s*^−1^ is a basal GTPase conversion rate, *γ* = 1 is its maximal rate and *K* = 1 is a saturation parameter.

The polarization process of GTPase is activated by an external stimulus that up-regulates the active form of GTPases that converts inactive into active forms. To model this external stimulus, we use a function *f*_*S*_(*ρ*^*S*^) = *k*_*S*_(*x, t*)*ρ*^*S*^, where *k*_*S*_(*x, t*) is the increased rate of conversion from inactive to active forms due to the external signal. Different forms of *k*_*S*_ are specified in next sections for specific cases of cell motility.

### 2.4 Mechanical model of the actin network

To model mechanically the actomyosin network, we consider an active, viscous segment in contact with the ECM [109, 102, 34]. We assume that viscous forces dominate the elastic forces and that inertial forces are negligible [90]. In this framework, the balance of linear momentum for the actomyosin network is

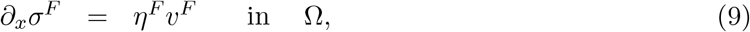

where *v*^*F*^ is the velocity of the retrograde flow. The r.h.s term represents the friction between the actin flow and the extracellular space, with friction parameter *η*^*F*^. The constitutive relation of the F-actin network stress is

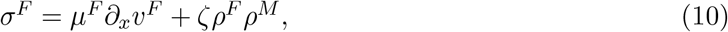

that accounts for the viscosity of the actin network and the myosin contractility. *µ*^*F*^ is the shear viscosity, *ζ* the active contraction exerted by the myosin motors and *ρ*^*M*^ and *ρ*^*F*^ are the density of myosin motors and F-actin, respectively, as we describe above.

We impose Neumann boundary conditions for the actomyosin network,

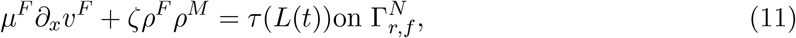

to compute the resulting velocity field and, consequently, the migration velocity (see Section 2.5 for details). We can consider three different situations. We can assume that the stresses at the boundaries of the actomyosin are zero, *τ* (*L*(*t*)) = 0 and, therefore, the actomyosin network flows by the only action of the myosin motors. We can also consider that an initial contraction of the cell, due to the myosin motor activity, or expansion, due to the actin polymerization at the front of the cell, will either decrease or increase the cell membrane length, respectively. These variations in cell membrane length induce a change in the membrane stress [37, 49] which results in a stress imposed on the actin network. To model the membrane tension, and the stress imposed in the network, we use a simple Hookean spring such that *τ* (*L*(*t*)) = *k*(*L*(*t*) *− L*_*b*_), where *k* is a spring constant and *L*_*b*_ = *L*_0_ + *L*_*r*_ accounts for the resting length, *L*_0_, and the buffer membrane length *L*_*r*_ in reservoirs and foldings of the cell membrane. Therefore, membrane tension starts to increase once all reservoirs and membrane folds have flatten, i.e *L*(*t*) *> L*_*b*_. In compression, *L*(*t*) *< L*_*b*_, we also use a simple linear element with constant spring constant 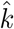. We analyze the physical implication and results of each of these BCs in following sections.

### 2.5 Effect of membrane tension in protrusion velocity and moving velocity of the cell membrane

Finally, we model the velocity of actin filaments polymerization against the cell membrane, *v*_*p*_, and the total protruding velocity of the membrane. Actin protrusion is regulated by the opposing membrane tension, the actin monomers availability anda mount of promoting factors of the actin nucleation.

We follow here a previous model of actin filament growth [103]. Filaments grow freely with velocity 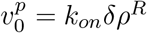 if there is no opposing force to actin polymerization. *k*_*on*_ is the rate of actin assembly, *δ* is the size of one single monomer at the tip of the filament and *ρ*^*R*^ is the density of GTPases that promote actin polymerization. We assume here that there is always a large pool of actin monomers available for polymerization, as we discussed in Section 2.1. Moreover, the actin network polymerizes and pushes against the cell membrane, which further builds membrane tension [103, 110]. Membrane tension has been characterized experimentally and theoretically in the past [111, 48, 112]. The membrane tension imposes a reaction force on the protruding actin network and if the membrane tension increases, the actin polymerization decreases [12, 13]. The actual velocity of actin polymerization at the cell membrane has been shown to decrease with respect to the free polymerization velocity as [113]

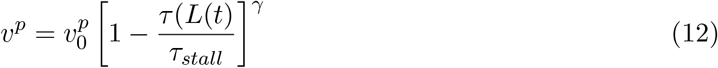

where *τ*_*stall*_ is a tension required to stall the actin network and *γ* is a model parameter that controls the velocity decay, which has been shown to be equal to 8 in keratocytes [113].

Finally, the protruding velocity of both sides of the cell is 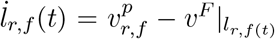 so that the outward polymerization velocity at the cell membrane, 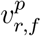, competes with the inward retrograde flow velocity *v*^*F*^ |_*r,f* (*t*)_ to expand or retract the cell boundary.

### 2.6 Numerical solution of the system of equations and model parameters

We solve computationally the system of Eqs. 1, 5, 6 and 9 in a staggered approach. We use the finite element method to discretize the system of equations in space [114] and an implicit-second order Crank-Nicholson method to discretize the parabolic equations in time [115, 116]. The complete finite element and time discretization procedure is presented in A. We use finite elements of size *h* = *L*(*t*)*/N*, where N is the number of elements.

The numerical solution of the parabolic equations in Section 2.1 and 2.2 can present undesired oscillations if the problem becomes convective dominant, i.e. *Pe >* 1, where *Pe* = *hv*^*F*^ */*2*D*_*F,M*_. As the convective velocity is the solution of Eq. 9, we cannot guarantee a priori that the problem will remain in the limit case of *Pe <* 1. Because we want to keep the number of elements of our domain constant and avoid remeshing strategies, we include a Stream-Upwind Petrov Galerkin (SUPG) stabilization term to overcome numerical oscillations in our solution [115, 116]. The numerical implementation has been previously tested against experimental and theoretical results [105].

To compute all simulation cases in following sections, we use one single set of model parameters from now on (see B, Table 1, for details) unless specified otherwise and only combine the different models described in previous sections.

## 3 Results

### 3.1 Early symmetric cell spreading

First, we focus on the early phase of symmetric cell spreading and provide a mechanistic rationale of the underlying forces (see Fig. 2). *l*_*r*_(*t*) and *l*_*f*_ (*t*) now represent the left and right sides of the cell, as the rear and front of the cell are not yet established. To reproduce the symmetric spreading, we consider that there is not polarization of signaling cues, i.e *ρ*^*R*^ = 1 and, therefore, the free polymerization velocity is 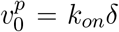. From now on, we use the transport models of one single form of actin and myosin (see Sections 2.1 and 2.2) because it allows us to compute the F-actin and bound myosin individually while reducing the computational cost of the simulation. The other modelling approaches, that is two-species for the actin and myosin forms and one single actomyosin phase, show minor differences in the model predictions (see C for details). We also impose homogeneous natural boundary conditions in the mechanics of the F-actin network (Eq. 36), i.e. zero stress. The effect of imposing a stress value as is an increase in the retrograde flow (see Fig. 12). We believe that zero stress BC provides a more realistic condition in the retrograde flow. We simulate the model predictions in time until steady state (no changes in the cell length).

**Figure 2:**
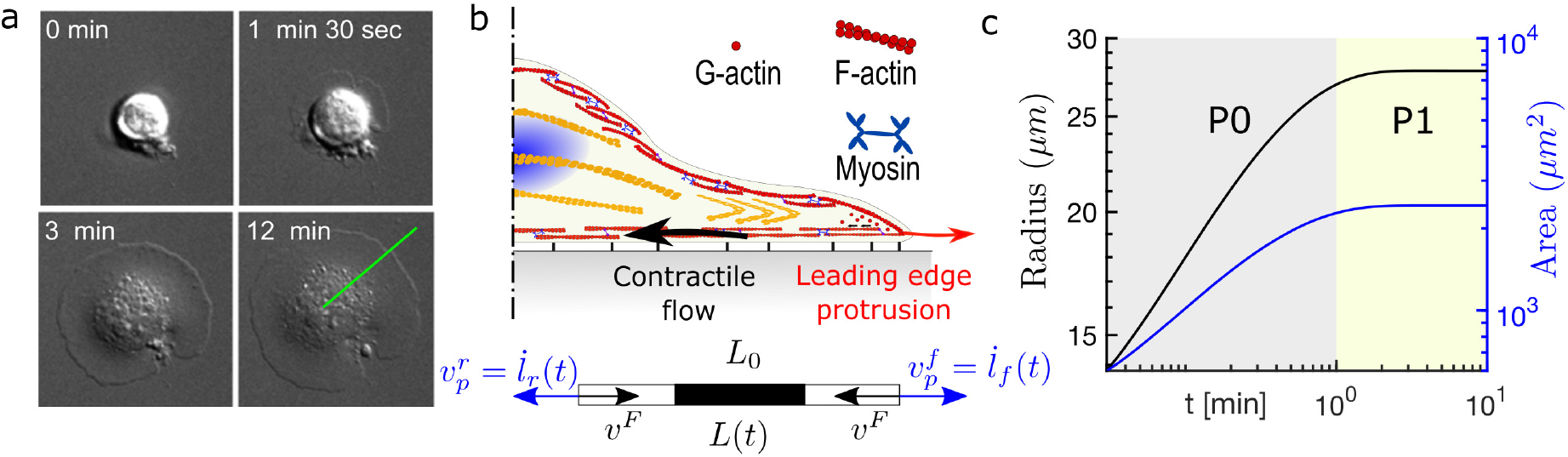
(a) Cell spreading process in time. Images adapted from [39]. (b) Sketch of the main forces acting in the cell spreading process: An active and contractile acto-myosin flow (black) and protrusive forces at the leading edge of the cell (red). At the cell-ECM interface, a friction-like force appears as a result of adhesion mechanisms. (c) Radius and area evolution of the spreading process. Results reproduce previous experimental data [117, 39]. Grey region shows the time period in which P0 takes place, up to ∼ 3 min. Yellow region shows the time period in which P1 occurs. P1 spans from ∼ 3 to 10 min.

First, we analyze the cell contact area and radius (Fig. 2c), which show a first phase of fast spreading that lasts ∼3min, where the radius of the cell increases ∼3-fold (P0). Then, the spreading rate decreases until steady-state, which occurs at ∼10min (P1). These results closely recapitulate previous results on the kinetics of cell spreading [118, 117, 39, 119, 47], where the initial stages of cell spreading show systematically a power-law for the cell radius.

We then analyze the evolution of the driving forces behind cell spreading. We show that the densities of actin and myosin at both sides of the cell are equal, which indicates a symmetric distribution of the intracellular components (Fig. 3a). The protrusion velocity of actin flow at the cell fronts starts at the free velocity, evolves symmetrically (Fig. 3b), and it takes its maximum value at the initial phase of spreading because there is no membrane tension resisting the F-actin polymerization at the cell front (Fig. 3c). A quick increase in membrane tension (Fig. 3c) up to 0.05 *nN/µm*) decreases the actin polymerization velocity at the cell front (Fig. 3b). Note that the membrane tension is in close agreement with previous experimental results [120]. The polymerization and retrograde velocities equal at ∼3 min (Fig. 3b). Once these two velocities equal, the spreading process reaches the steady state, i.e. 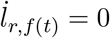. The results also indicate that the quick spreading of the cell restricts the acto-myosin network to the center of the cell during the first 3 min of the spreading process. The accumulation of contractile myosin motors in the cell center is responsible for activation and acceleration of the retrograde flow (Fig. 3b). Then, the myosin and actin concentrations evolve to a less but still polarized state because of diffussion and turnover. We also show the kymographs of the actin and myosin densities as well as of the retrograde flow (Fig. 3d)

**Figure 3:**
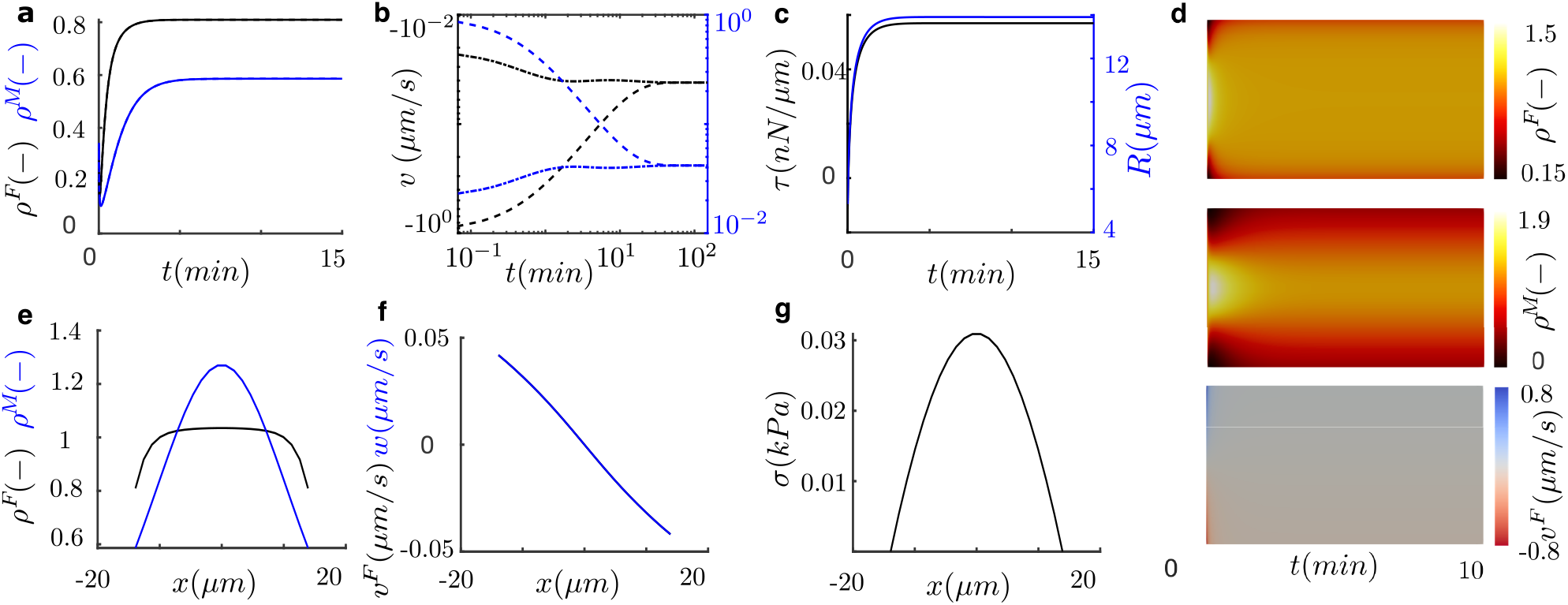
(a-d) Time and space evolution of cell spreading. (a) Actin (black) and myosin (blue) densities at the left (dash) and right (solid) fronts of the cell. Left and right variables are superimposed as a result of their symmetric distributions. (b) Polymerization velocity (dot-dash) and retrograde velocity at the cell membrane (dash). (c) Membrane tension (black) and cell radius (blue). (d) Kymographs of the actin density (top), myosin density (center) and retrograde flow velocity (bottom). (e) Actin (black) and myosin (blue) densities, (f) retrograde flow at the cell (blue) and lab frame (black) and (g) tension of the actin network at steady state along the cell length.

At steady state, we show a symmetric distribution of actin and myosin (Fig. 3e), with an accumulation of actin and myosin at the cell centre of ∼1.08 and 1.5, respectively. We also see a symmetric retrograde flow from both sides toward the cell center, where it vanishes (Fig. 3f). The maximum actin flow velocity is *v*_*F*_ = 0.045*μm/s* at the cell sides, which is again in agreement with previous data [29, 91, 101]. Consequently, the stress of the actomyosin network is also symmetric with a maximum at the cell center of 30Pa (Fig. 3g). This result indicates that the stress of the actomyosin network imposes a traction on the nuclear region of the cell, which is in agreement with previous results on the mechanosensitivity of the cell nucleus [121].

Our results confirm that the dynamics of the cell spreading is controlled by the retrograde flow and the actin polymerization at the front of the cell. While the actomyosin network flows inward, powered by myosin motors at the cell center, the protrusive actin moves the membrane forward. The actual cell spreading velocity is the result of these two competing flows. Our results confirm previous experimental data showing that the decrease of the initial spreading velocity is not only due to the reduction of the actin protrusion velocity but also due to the effect of the inward retrograde flow [122, 39]. Our simulations show that the P1 phase of cell spreading is distinguished by a high protrusion velocity and an increasing retrograde flow. The slow retrograde flow and the buffering of the cell membrane reservoirs are responsible for the fast protrusion of the cell membrane during the first seconds of the process. When sufficient membrane tension is reached, the actin polymerization velocity decreases which, together with the decrease in the retrograde velocity, causes the transition from P0 to P1. Eventually, the opposing actin networks equals and the spreading process reach the steady state.

We also validate our model of symmetric cell spreading by reproducing the effect of substrate rigidity [123, 124, 125] and myosin contractility inhibition [122] (see C and Section 1.2). Our simulations agree with previous results that showed an increase in cell spreading as the substrate rigidity increase (C). This is due to a weaker retrograde flow that competes with the free velocity of the actin polymerization. Similarly, a diminished retrograde flow induced by the inhibition of the myosin contractility is responsible for the increase of spreading area. The impaired inward retrograde flow allows the cell front to protrude faster, with a spreading velocity closer to the free polymerization velocity (Fig. 14, C).

### 3.2 Cell migration

#### 3.2.1 Mesenchymal cell migration

We start the migration study analyzing mesenchymal cell migration. To break the symmetric spreading, we polarize the signaling cues described in Section 2.3. We introduce a random stimulus, *f*_*S*_(*ρ*^*S*^), and let the densities of active, *ρ*^*R*^, and inactive, *ρ*^*S*^, GTPases to evolve in time (see Fig. 15). We associate the active GTPases, *ρ*^*R*^ to the GTPase exchange factor (GEF) [53]. We assume that an increase of GEF instantaneously activates protrusion-related mechanisms controlled by Cdc42 and Rac1 and the polymerization velocity following Eq. 12. An increase in GEF is associated with decrease in the downstream alterations in GTP-RhoA, membrane exocytosis and myosin activation, but it is promoted where GEF is reduced [53, 51, 59, 10]. Because Rho and Rac1 are mutually exclusive [65, 66], we use the mirror of 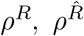, as an indicator of GTP-RhoA and modify the constitutive relation of the F-actin mechanics (Eq. 10) as 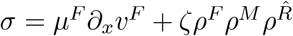 to activate the myosin contractility just in regions where GTP-RhoA is accumulated.

As in the symmetric cell spreading case, cell migration is promoted by the competition of the actin polymerization against the cell membrane in the cell periphery and the inward retrograde flow. During migration, these motile forces are polarized front-to-back and, in consequence, make the cells to migrate in a specific direction. The actin and myosin densities accumulates in the back of the cell (Fig. 4a,e) because an asymmetric retrograde flow drags them backward (Fig. 4b,f). The retrograde flow is activated by the effect of the GTP-RhoA polarization, 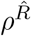, which induces myosin contractility in rear of the cell (Fig. 4d,f,h). The actin network flows inward with *v*_*F*_ ≈ 0.1*μm/s* at the front and rear, although there is a larger portion of the cell with a positive velocity, which explains the accumulation of densities at the right of the cell domain. The free polymerization velocity is now 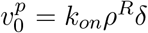 (Fig. 4b) and, therefore, it increases at the cell front, where *ρ*^*R*^ accumulates (Fig. 4d,h) and decreases due to the increase in membrane tension (Fig. 4c). We also show the space-time evolution of the densities and retrograde flow in a kymograph (Fig. 4i). Our results indicate that there must be an upstream signaling polarization that modifies the motile forces of the cell to break the symmetric spreading. Our model shows that the effect of an increased polymerization of actin and activity of myosin motors at the cell front and rear of the cell, respectively, enable directed cell migration. These results just confirm many previous data on cell migration, from the need of a polarized signal [10], to the downstream adaptation of the actin polymerization and the myosin activity [50, 10, 51, 52].

**Figure 4:**
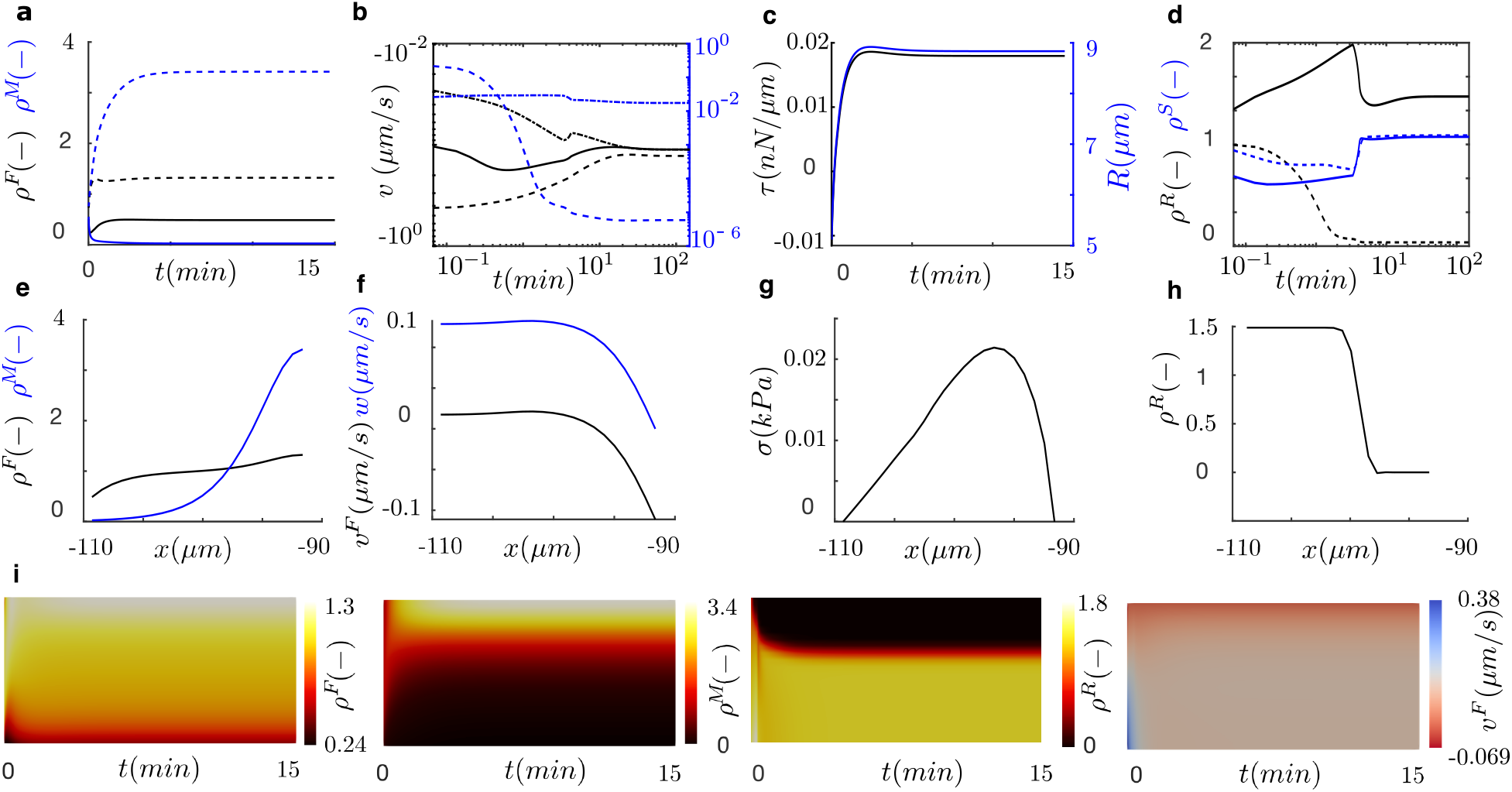
Mesenchymal cell migration. Time evolution of (a) actin (black) and myosin (blue) densities at the center (dash) and front (solid) of the cell, (b) polymerization velocity (dot-dash), retrograde velocity at the cell membrane (dash) and total velocity of protrusion (solid), (c) membrane tension and (d) *ρ*^*R*^ (black) and *ρ*^*S*^ blue in the front (solid) and rear (dash) of the cell. At steady state, (e) actin (black) and myosin (blue) densities, (f) retrograde flow in the lab (black) and cell (blue) frame, (g) tension of the actin network and (h) density of the active signals along the cell length. (i) Kymographs of the actin density, myosin density, *ρ*^*R*^ and retrograde flow velocity.

Finally, we wonder whether actin polymerization at the cell front is has a stronger or weaker effect on the migration velocity than the contractile actomyosin flow. We inhibit the effect of Rac1 in actin polymerization and of RhoA in myosin motor activity (Fig. 5). The results indicate that actin polymerization at the cell front has a stronger effect in cell migration than the myosin activity. Details on other model results are given in Fig. 16 and Fig. 17.

**Figure 5:**
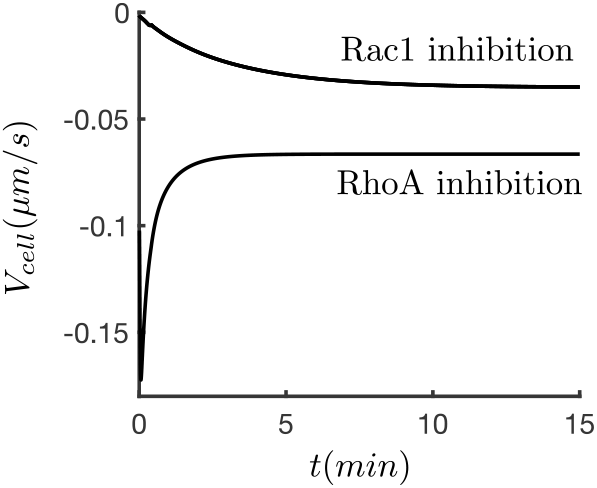
Inhibition of Rac1 in actin polymerization and of RhoA in myosin motor activity. Cells with an inhibited effect of Rac1 on actin polymerization migrate with *V*_*cell*_ ≈35*nm/s*. Cells with an inhibited effect of RhoA on myosin activity migrate with *V*_*cell*_ ≈ 65*nm/s*.

#### 3.2.2 Ameboid cell migration

Next, we ask ourselves if the same model of mesenchymal migration may reproduce and explain the case of ameboid cell migration. Ameboid cell migration is expressed in confined environments such as those found in 3D porous tissues [69, 71, 67] (see Fig. 6 and Section 1.4 for details). In this type of cell migration, the actin polymerization against the cell membrane is manifested weakly, i.e. *v*_*p*_ = 0. To tackle this mode of migration, we impose Neumann BC as we described in Section 2.4, which induces an outward actin flow in the lab frame of reference that controls directional migration [126]. This outward velocity at the cell front accounts for the protrusion of the cell front, in form of blebs, and the contraction of the cell rear, that are responsible for the cell translocation [69, 71, 67]. The cell initially contracts due to the action of the myosin motors because there is no actin polymerization that compensates the retrograde flow. Then, the membrane length decreases and the actin network experiences compressive forces.

**Figure 6:**
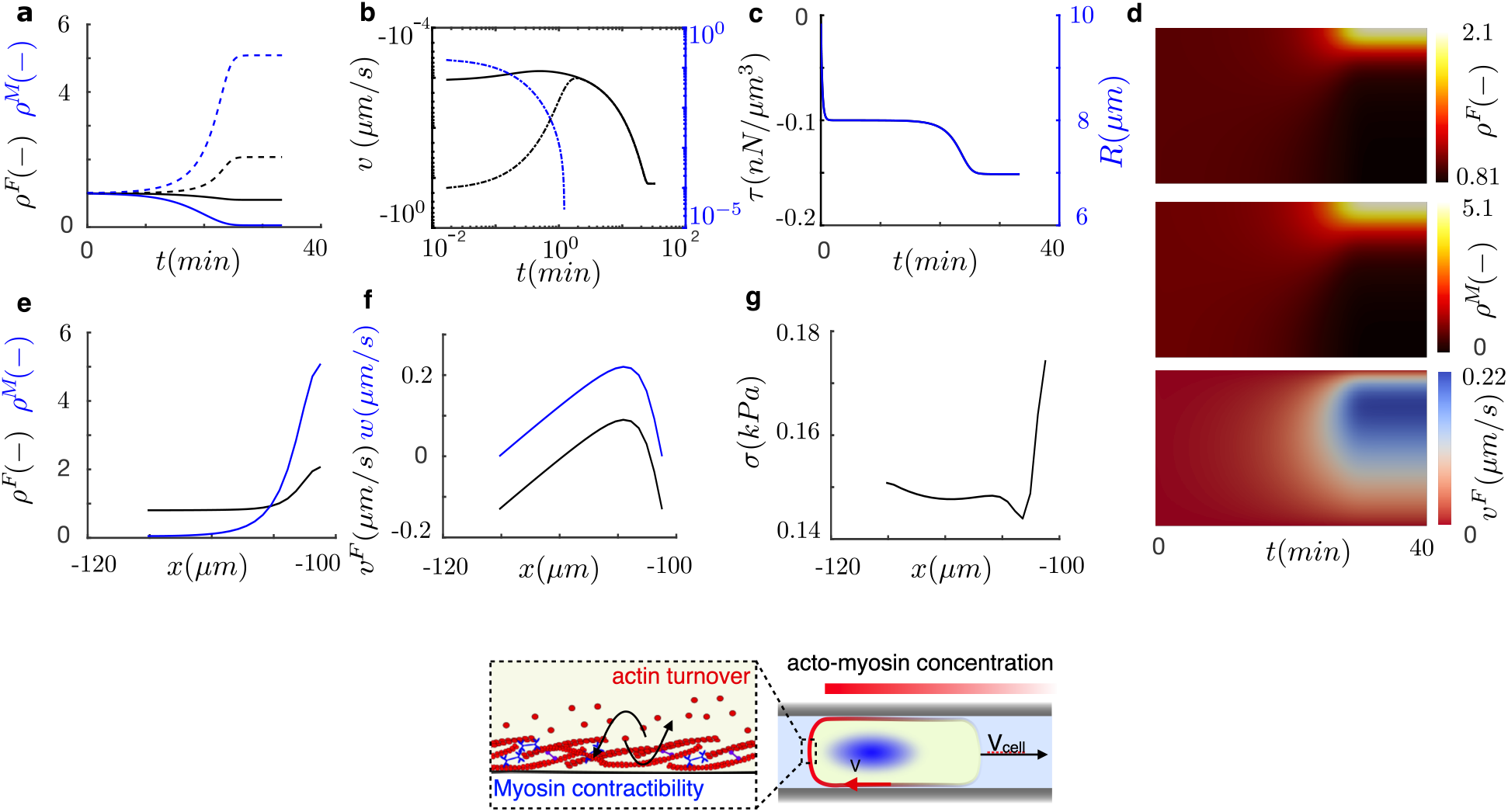
Ameboid cell migration. Time evolution of (a) actin (black) and myosin (blue) densities at the center (dash) and front (solid) of the cell, (b) polymerization velocity (dot-dash), retrograde velocity at the cell membrane (dash) and total velocity of protrusion (solid), (c) membrane tension (black) and cell radius (blue). (d) Kymographs of the actin density (top), myosin density (center) and retrograde flow velocity (bottom). At steady state, (e) actin (black) and myosin (blue) densities, (f) retrograde flow in the lab (black) and cell (blue) frame and tension of the actin network (g) at steady state along the cell length. Bottom, sketch of the ameboid mode of migration

During the first minutes, cells contract quickly (Fig. 6b,c) with an almost uniform distribution of actin and myosin motors, that is maintained during ≈5min (Fig. 6a). Then, the effective velocity of the cell increases due to the imposition of the stress boundary conditions, which results in a convective velocity that drags the actin network in the direction opposite to migration (Fig. 6b). This backward flow polarizes the acto-myosin concentration during ∼25min (Fig. 6a). The, the internal active forces of the crawling cell balance the frictional forces and the cell reaches a steady state migration velocity of 0.13*μm/s*, in agreement with previous data [69, 71]. At steady state, the densities of actin and myosin motors have increased 2- and 6-fold at the cell rear, respectively (Fig. 6e). The retrograde flow in the cell frame is maximum close to the cell rear and vanishes at the cell sides (Fig. 6f). These densities and retrograde flow velocities are also in close agreement with previous experimental data in confined cell migration [69]. The space and time evolution of those quantities are again shown in a kymograph (Fig. 6d).

Our full model is able to reproduce ameboid migration by just inhibiting the actin polymerization at the cell fronts and modifying the boundary condition to effectively translocate the cell body forward, similarly to previous models [126]. We show that imposing a stress value in the ameboid-like migration accelerates the retrograde flow, which polarised the actomyosin network and, eventually, the migration velocity. Therefore, the stress BC may represent a plausible case to model the protruding velocity of the membrane in ameboid-like migration. Our results indicate that the cells do not need a polarization of signals because the system self-polarizes when the retrograde flow is induced by the stresses imposed in the network.

### 3.3 Tug of war between taxis

In section 3.2, we have validated our full model of cell motility in different phases and modes of cell migration against previous theoretical and experimental data. We have used one single model, with one set of material parameters to reproduce all of them. Now, we use the mesenchymal cell migration model to analyze the effect of chemical and mechanical cues in the directing cell migration. Then we study how the motile mechanisms of the cell competes with each other when chemical and mechanical cues appear simultaneously, which has not been yet investigated neither experimentally nor theoretically.

#### 3.3.1 Chemotaxis

We start the chemotactic analysis with a polarized cell that moves in one direction, as we described in Section 3.2.1. Once the cell is close to its steady state, at ≈ 8min we induce a chemical stimulus in the direction opposite to the current migration. We see that such a stimulus is capable of switching the polarization of the GTPases in the cell, which reaches its steady state in less than 3 minutes (see Fig. 7d,i and Fig. 18).

**Figure 7:**
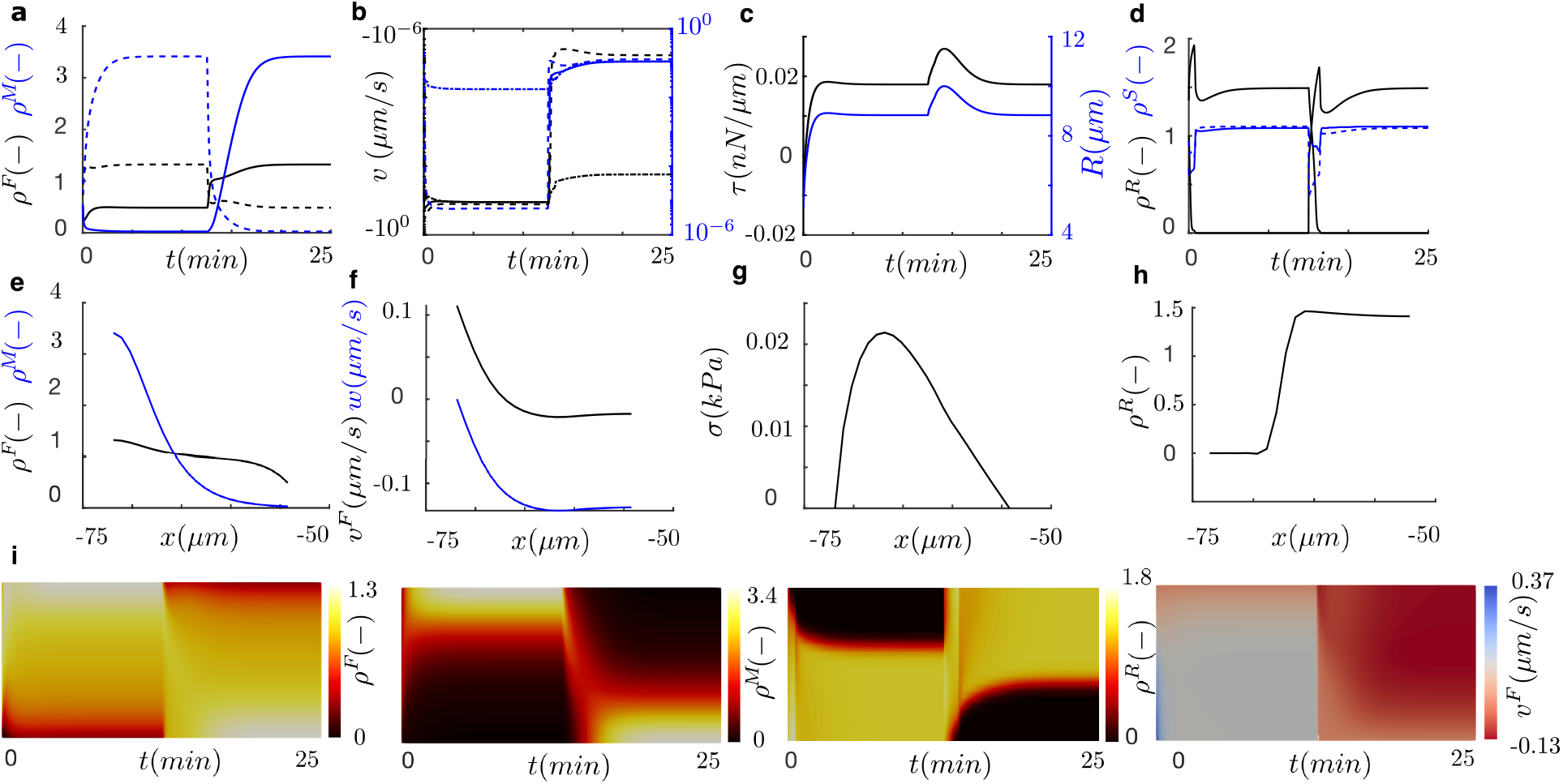
Chemotaxis in mesenchymal cell migration. Time evolution of (a) actin (black) and myosin (blue) densities at the center (dash) and front (solid) of the cell, (b) polymerization velocity (dot-dash), retrograde velocity at the cell membrane (dash) and total velocity of protrusion (solid), (c) membrane tension (black) and cell radius (blue) and (d) *ρ*^*R*^ (black) and *ρ*^*S*^ (blue). At steady state, (e)actin (black) and myosin (blue) densities, (f) retrograde flow in the lab (black) and cell (blue) frame and (g) tension of the actin network along the cell length. (i) Kymographs of the actin density, myosin density, *ρ*^*R*^ and retrograde flow velocity.

Once the GTPases polarize, the downstream motile mechanisms of the cell also polarize. The front moves from one side to the other of the cell (Fig. 7b,i) because of the repolarization of GTPases involved in actin polymerization (Fig. 7d,i). Simultaneously, the activation of myosin motors, due to the accumulation of RhoA, also shifts from one side to the other of the cell, which also repolarizes the direction of the retrograde flow (Fig. 7a,b,i). There is also a sudden increase in the cell length and, consequently, in membrane tension during the repolarization phase (Fig. 7c). During this transient phase the polymerization velocity and the retrograde velocities equals at both ends, which resemble the quick symmetric spreading mode. At steady state, the density and velocity profiles are the same as the ones reported in Section 3.2.1 but with the front-back polarization shifted (Fig. 7e,f). The kymograph clearly shows the shift in the intracellular variables (Fig. 7i). The cell migrates with the same velocity as before the chemotactic stimulus but in opposite direction (Fig. 7). In summary, the model predicts that a chemical stimulus is capable of polarazing cellular signals, thus completely shifting the migration direction of the cell, as it has been demonstrated experimentally before [72, 73, 127].

#### 3.3.2 Durotaxis

Then, we analyze the case of single cell durotaxis. To analyze in isolation the effect of a ECM stiffness gradient in cell motility, we cancel the sudden polarization of GTPases so that they cannot establish the direction of migration. Therefore, we start with the symmetric cell spreading case described in Section 3.1. To reproduce the effect of a stiffness gradient in the ECM, we take traction forces, *P*, and actin flow velocities, *v*, from previous experimental data as a function of the ECM stiffness (E *∈*0.1-100 kPa)[128] (Fig. 8a) and compute the effective friction parameter as *η* = *P/v*. To change the stiffness gradient of the substrate, we define samples of varying length (100, 200 and 2000 *µ*m) with the same stiffness range (E*∈*0.1-100 kPa). We also consider three initial locations at the sides and centre of the sample where cells are initially seeded to study how the absolute value of the substrate stiffness, and not only its gradient, may regulate the durotactic response.

**Figure 8:**
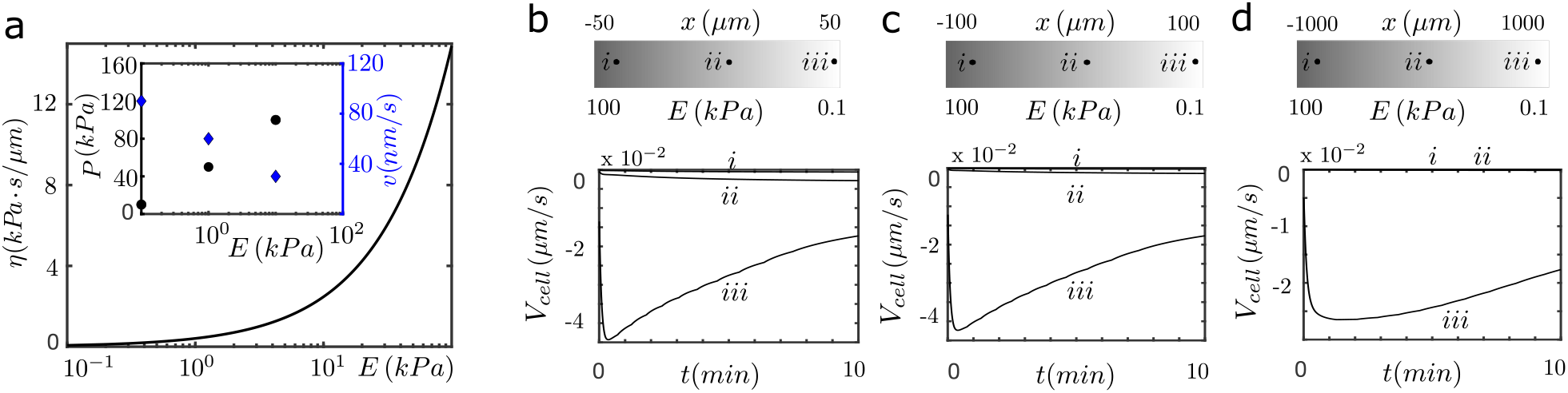
(a) Friction parameter prediction as a function of the substrate rigidity from traction and retrograde flow velocity experimental data []. (b-d) Cell migration velocity in different durotactic environments. The substrate stiffness goes from 100 kPa, at the right of the sample, to 0.1 kPa, at the left. Points i, ii and iii represent the locations where the cells are initially seeded. (b) Sample of 100*μm* in length, i=-30 *µm*, ii=0*μm*, iii=30 *µm*. (c) Sample of 200 *µm* in length, i=-80 *µm*, ii=0 *µm*, iii=80 *µm*. (d) Sample of 2000 *µm* in length, i=-845 *µm*, ii=0 *µm*, iii=845 *µm*.

The results of our model reproduce the well known fact of migration toward positive gradients of substrate rigidity (negative velocity in all our results, Fig. 8) found in most cell types and, specifically, in the ones from where the friction values we used here are obtained [128]. The migration velocity is maximum in cells moving from the softest location in each sample (Fig. 8b-d, points *iii*) with *V*_*cell*_ ≈ 25 − 45*nm/s*. For cells initiating the migration in this soft location, the maximum migration velocity is obtained in cells moving in the largest gradient, that is in samples of L=100 *µm* (Fig. 8b, point *iii*), where *V*_*cell*_ = 45*nm/s*. However, cells seeded on large stiffness substrates (points *i, ii*, Fig. 8b-d) do not undergo durotaxis in any of the samples and they only sustain an almost symmetric spreading. These results indicate that cell migration is favoured in soft matrices with large friction gradients. This is because large stiffness gradients induce a substantial polarization of the actomyosin network (Fig. 20-21) which enables the retrograde flow at the cell rear to translocate the cell forward. In all the simulations, the cells slow down as the cells migrate toward the stiffer regions (see results at for longer simulation time Fig. 19). This is particularly noticeable on cell initially seeded in soft locations. For reference, we show the evolution of all model variables in space and time for the most prominent durotactic case (larger stiffness gradient, L=100 *µm*) for cells seeded at the softest regions (point *iii*) and at the stiffest location (point *i*) in Fig. 20 and 21, respectively. We also show the model variable at the lowest stiffness gradient (L=2000 *µm*) and at soft locations (point *iii*) in Fig. 22, which, as we discussed above, present a symmetric distribution similar to the spreading case.

#### 3.3.3 Competition between chemotaxis and durotaxis

Finally, we investigate the effect of imposing simultaneously chemical and mechanical cues in the motile cell. Because both signals can be simultaneously present in vivo, we aim to understand what signal, chemical or mechanical, has a stronger effect in persistent cell migration. To do so, we place again cells in the same substrates of varying gradient stiffnesses and lengths that we describe above. We activate GTPases signaling at initial time to reproduce an imposed or natural exogenous chemical signal, as explained in Section 3.2.1. We simulate a chemotactic signal that opposes to the stiffness gradient and another where the chemical signal is oriented in the same direction as the durotactic cue.

The chemical signal quickly activates the GTPases and the downstream actomyosin polarization, as we describe in Section 3.2.1. This is a fast process that establishes the front and rear of the cell in few minutes (Fig. 18). The cell starts to migrate in the direction of the chemical stimulus (Fig. 9) because the protrusive actin polymerization pushes the cell in the direction of the formed front while the rear retrograde flow translocates the cell rear forward.

**Figure 9:**
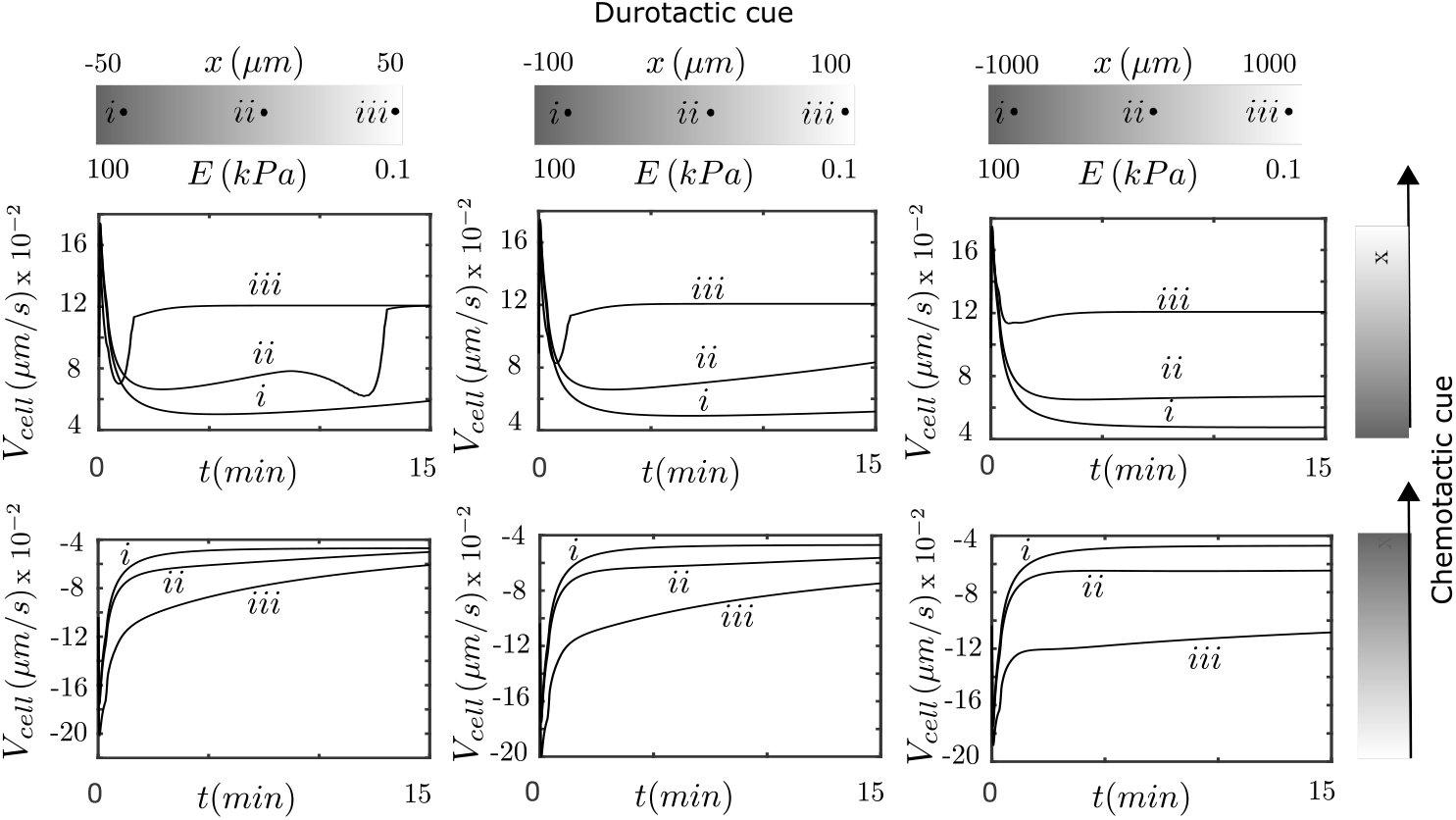
Cell migration velocity under simultaneous durotactic and chemotactic environments. Organized by column, samples of 100, 200 and 2000 *µ*m length, from left to right. The substrate stiffness goes from 100kPa, at the right of the sample, to 0.1 kPa, at the left. Points i, ii and ii represent the location where the cells are initially seeded. In first column, samples of 100*μm* in length, i=-30*μm*, ii=0*μm*, iii=30*μm*. In second column, samples of 200*μm* in length, i=-80*μm*, ii=0*μm*, iii=80*μm*. In the thrid column, samples of 2000*μm* in length, i=-845*μm*, ii=0*μm*, iii=845*μm*. Organized by row, two chemical stimuli in opposing directions. Results in the first row show a chemical cue in the same direction as the durotactic cue. Results in the second row represent cases where a chemical cue is set in opposite direction to the durotactic cue.

When both signals are oriented in the same direction, the migration velocities increases with respect to the durotactic signal alone for all simulated cases (Fig. 9, bottom). Indeed, the total migration velocity is approximately the sum of the single durotactic and chemotactic migrations (see Fig. 7 and Fig. 4). Therefore, the chemical and mechanical stimuli induce a cooperative response in the motility forces of the cell. When the signals are set in opposite directions, we see different responses (Fig. 9, top). When cells are placed at stiff parts of the substrate, the chemotactic cue rules over durotaxis, and cells migrate guided by the chemical signals, although the opposing durotaxis progressively slows down the migration velocity. When the migrating cells are located at the lowest substrate rigidity (Fig. 9, iii), the chemical signal also dominates the directed migration at the beginning but the large stiffness gradient quickly slows down the migration velocity. This is because, as we explain above, the polarization of the retrograde flow due to the durotactic response is maximized in this specific location. The constant migration velocity after ≈3min is because, once the cell reaches the limit of the sample, we keep the substrate rigidity constant. In summary, a combination of a large substrate stiffness gradient in soft matrices may inhibit chemotaxis.

All these differences occur due to the competition of the polymerization velocity and the retrograde flow at the front and rear of the cell. We show the evolution of all model variables in space and time for the softest location (point *iii*) and largest stiffness gradient (L=100*μm*), in aligned (Fig. 25) and opposed taxis (Fig. 26). As before, the and also at the stiffest location (point *i*) for aligned (Fig. 23 and Fig. 24). We also show the model variables at the lowest stiffness gradient (L=2000*μm*) at the softest location (point *iii*) in Fig. 27 and Fig. 28) for aligned and opposite cues. Despite the actin flow polarization in large friction gradients in soft matrices, the asymmetric concentration of GTPases induces a larger actin polymerization in what it becomes the cell front, and a large myosin contractility in the opposite side of the cell, which directly competes with the polarization induced by the stiffness gradient. When the durotaxis induces a strong polarization of the actomyosin network, it reduces the effect of chemotaxis but cannot direct cell migration.

Finally, we wonder how we could effectively inhibit the exogenous chemotactic cues so that we could control the directing migration through durotaxis. Because small chemical signals are often present in in-vivo and in-vitro systems, it is important to understand how they may interact and compete with stiffness gradients of the ECM. As we show in Section 3.2, inhibiting actin polymerization at the cell front and myosin activity may arrest cell migration. We hypothesize that canceling any of these two motile forces may inhibit the chemotactic direction. To do so, we first cancel the dependance of the polymerization velocity on the GTPases signal *ρ*^*R*^. As a consequence, the polymerization velocity at the rear and front of the cell are equal and, therefore, only depend on the cell membrane tension. We still allow the GTPases to activate myosin contractility at the rear of the cell. Second, we inhibit the dependance of myosin contractility on GTPases but allow the polymerization velocity to depend on the polarized signal *ρ*^*R*^. Then we simulate again cell migration under simultaneous durotactic and chemotactic signals (Fig. 10). As in the control case of the tug-of-war between chemotaxis and durotaxis, the alignment of both signals in the same direction increases the velocity of migration with respect to the pure durotactic signal. However, chemotaxis is now weakly expressed and, therefore, the migration velocities are reduced with respect to the control tug-of-war between chemotaxis and durotaxis. When signals are opposed to each other, cells placed in the soft part of the sample undergo durotaxis although the chemical signal quickly slows down the migration velocity. On the other hand, cells placed in intermediate and high stiffness regions follow the chemical signal although the migration velocity is lower than in the control case. Our results show that durotaxis may rule over chemotaxis in soft substrates but chemotaxis controls migration directionality in stiff substrates. For comparison purposes, we show all model parameters for the L=100*μ*m length sample and cells seeded at the softest location for aligned and opposite cues in Fig. 29 and 30, respectively.

## 4 Conclusions

For decades, theoretical and computational models have provided a fundamental understanding of the underlying forces that enable cell motility. Some previous models have focused on specific mechanisms of cell motility, such as the actin polymerization velocity [103, 13] or the mechanics of the retrograde flow [109, 34]. Others studied particular modes of cell motility, including cell spreading [129, 118, 130], cell migration [101, 79] and individual cell taxis [74, 94, 77, 95].

In this work, we developed a complete mechanical model for cell motility that integrates the most important mechanisms involved in cell motility from previous theoretical and computational models. Specifically, we put together the signaling of molecular cues, the mechanics and dynamic distribution of the actin network and the contractile forces of the myosin motors, which together represent the actomyosin network. By integrating the contractile actomyosin network adhering to the ECM [103, 90, 101, 102] with a network of actin that polymerizes against the cell membrane [103, 113], we were able to model the most important phases of cell migration and distinctive modes of cell migration through one single model of cell motility, using also one single set of model parameters. This is an important aspect from a modeling point of view because it allows to pinpoint the forces that controls cell motility across modes and phases. We validated this full model of cell motility against well know results of symmetric spreading [39, 131, 9], confined cell migration [69] and polarization-driven ameboid-like migration [79, 101]. The sole minimalistic model used in this study has not only been able to reproduce all these previous experimental findings, but also to describe the universal underlying physical forces that control cell motility.

These scenarios may vary across cell types, depending on the effect of, e.g, GTPases in actin polymerization or myosin activity, in the adhesion mechanics, specific turnover rates of actin, or the mechanics of the cytoskeleton. Variations in all these mechanisms may explain discrepancies in migrating behaviors across cell types. Also, other mechanisms may be involved in migration modes. For example, it was shown that confined cells may migrate by just balancing fluxes of water across the cell, which allows the front and rear of the cell to translocate the cell body forward. This water-based mechanism of cell motility has been effectively recovered by the mathematical models [71, 132].

We then used our validated model to analyze chemotaxis and durotaxis in isolation and in competition. We showed that a small and sudden chemical stimulus was capable of polarising the cell and of imposing a directional migration. We also showed that cells need a large stiffness gradient on soft substrates in order to enable a durotactic response. Indeed, cells placed in these same soft locations show increasing motility for increasing stiffness gradients. Cells seeded in stiff substates showed slow migration or almost symmetric spreadings. There are still important opened questions. For example, neurons show persistent directionality toward softer rigidities [3], while epithelial tissues travel to stiffer regions [74]. So how different cell types respond to stiffness gradients in opposing directions?

More importantly, we use our model to investigate a tug-of-war between chemotaxis and durotaxis. The understanding of a tug-of-war between durotaxis and exogenous chemical signals that appears in in-vitro and in-vivo experiments is key in tissue engineering and to design better biomimetic materials. Although chemotaxis and durotaxis has been investigated before in isolation, whether durotaxis overrules chemotaxis, or vice-versa, and how the motile forces of the cell compete in this situation has been overlooked. We show that chemotaxis controls cell migration in all the situations analyzed. When the chemical and mechanical signals cooperate migration is enhanced. When chemical signals and stiffness gradients are imposed in opposing directions, chemotaxis can strongly diminished when cells are seeded in strong durotactic locations, that is soft substrates with large stiffness gradients, but still controls the directed migration. However, we showed that inhibiting the effect of GTPases in the motile forces of the cell, specifically in the protruding polymerization of actin, make durotaxis prevalent in soft matrices.

In summary, we have reproduced the main phases and modes of cell migration, including chemotaxis and durotaxis. Moreover we showed how chemical and mechanical cues compete to each other. These mechanisms and modes of cell motility are analysed every day in thousands of labs around the world to study, for example, why some cells follow positive stiffness gradient while others follow negative durotaxis [74, 3] or how cell invasion can be arrested [133]. To assist in the answer of these and other questions in cell motility and in the design of optimized experiments, we provide a freely-available platform to analyze, by setting different model flags, results of each case discussed above (uploaded in *to be uploaded upon acceptance, it includes personal information*). We believe that such tool, easy to set-up, may be used by experimental biologists in the community, to test hypothesis and rationalize observations, but also by mechanicians and biophysicists to further explore computationally cell motility.

## Acknowledgement

To be completed upon acceptance to ensure the double blind process.

