## appendices for "A full computational model of cell motility: Early spreading, cell migration and competing taxis"

September 28, 2022

### Appendices

#### A Time and spatial discretization

##### A.1 Transport equations

Let us consider a general density $\rho(x,t)$ and the parabolic equation that defines every transport equation involved in the models presented in Sections 2.1, 2.2 and 2.3

$$\begin{cases} \rho^t + (v\rho - D\rho_x)_x = s - \sigma\rho & \text{in } \Omega \times [0,T] \\ v\rho - D\rho^x = 0 & \text{on } \partial\Omega, \end{cases} \tag{13}$$

where $v$ is a convection velocity, $D$ a diffusion constant and $\sigma$ and s constants.

###### A.1.1 Time Discretization

We perform the time integration of our transport equations through the $\theta$ family method. Specifically, we use a second-order implicit, Crank-Nicolson method where $\theta = 1/2$,

$$\frac{\Delta(\bullet)}{\Delta t} - \frac{1}{2}\Delta(\bullet)_t = (\bullet)^n_t, \tag{14}$$

where $\Delta(\bullet) = (\bullet)^{n+1} - (\bullet)^n$ and $(\bullet)$ represents the concentration quantity of interest. We set a time interval $\mathcal{T}$ and a number of subintervals $n$, $\mathcal{T} = \bigcup_0^{n-1}[t^n, t^{n+1}]$, where the time increment is $\Delta t = t^{n+1} - t^n \geq 0$. The temporally discretized form of the general parabolic equations reads

$$\frac{\rho}{\Delta t} + \theta\mathcal{L}(\rho) = \frac{\rho^n}{\Delta t} - (1-\theta)\mathcal{L}(\rho^n) + s, \tag{15}$$

where $\mathcal{L}(\bullet) = \partial_x(v(\bullet) - (D\partial_x(\bullet)) + \sigma(\bullet)$ acts on $\rho$. Note that for sake of simplification the superindex for the time increment $n+1$ have been dropped.

###### A.1.2 Weak Form of the semi-discretized transport equation

We follow a standard Galerkin formulation for the weak form of the semi-discretized Eq. 15. We define a collection of trial functions, $\mathcal{S}_t$, that fulfill the Dirichlet condition in $\Gamma_D$, and a collection of test function, $\mathcal{V}$, that are square integrable, have square integrable first derivatives in $\Omega$ and vanish in $\Gamma_D$, i.e:

$$\begin{aligned} &\mathcal{S}_t = \{\rho | \rho(\cdot,t) \in \mathcal{H}^1(\Omega),\ t \in [0,T] \text{and } \rho(x,t) = \rho^d \text{ on } \Gamma_D\} \\ &\mathcal{V} = \{\omega \in \mathcal{H}^1(\Omega) \mid \omega = 0 \text{ on } \Gamma_D\} \end{aligned} \tag{16}$$

The weak form of transport problem in Eq. 15 takes the following form:
Find $\rho \in \mathcal{S}_t$, for any $t \in [0,T]$ such that

$$\int_\Omega \omega\frac{\rho}{\Delta t}\mathrm{d}\Omega + \theta\int_\Omega \omega\mathcal{L}(\rho)\mathrm{d}\Omega = \int_\Omega \omega\frac{\rho^n}{\Delta t}\mathrm{d}\Omega - (1-\theta)\int_\Omega \omega\mathcal{L}(\rho^n)\mathrm{d}\Omega + \int_\Omega \omega s\mathrm{d}\Omega \quad \forall\omega \in \mathcal{V} \tag{17}$$

Then, we apply Gauss theorem on the total flux term, considering the boundary such that $\partial\Omega = \Gamma_D \cup \Gamma_N$ and noting that $v\rho - D\rho^x = 0$. The problem reduces to find $\rho \in \mathcal{S}_t$, for any $t \in [0,T]$ such that

$$
\begin{aligned}
(\omega, \frac{\rho}{\Delta t}) + \theta[a(\omega,\rho) + c(v;\omega,\rho) + (\omega,\sigma\rho)] = \\
(\omega, \frac{\rho^n}{\Delta t}) - (1-\theta)[a(\omega,\rho^n) + c(v;\omega,\rho^n) + (\omega,\sigma\rho)) + (\omega, s)\ \forall \omega \in \mathcal{V}
\end{aligned}
\tag{18}
$$

where

$$
(\omega,\rho) = \int_\Omega \omega\rho \mathrm{d}\Omega, \qquad c(v;\omega,\rho) = \int_\Omega \partial_x \omega v \rho \mathrm{d}\Omega \quad \text{and} \qquad a(\omega,\rho) = \int_\Omega \partial_x \omega (D\partial_x \rho)\mathrm{d}\Omega \tag{19}
$$

Analogously to the spatial operator above, we define

$$
\hat{\mathcal{L}}(\bullet) = \int_\Omega \partial_x \omega v(\bullet)\mathrm{d}\Omega + \int_\Omega \partial_x \omega D \partial_x(\bullet)\mathrm{d}\Omega + \int_\Omega \omega\sigma(\bullet)\mathrm{d}\Omega \tag{20}
$$

so that we can rewrite the weak form in a more compact form:

$$
(\omega, \frac{\rho}{\Delta t}) + \theta\hat{\mathcal{L}}(\rho) = (\omega, \frac{\rho^n}{\Delta t}) - (1-\theta)\hat{\mathcal{L}}(\rho^n) + (\omega, s) \tag{21}
$$

##### A.1.3 Spatial Discretization

We then discretized in space the final semi-discrete weak form following a Galerkin finite element approximation. We define the finite dimensional subset of spaces $\mathcal{S}^h$ and $\mathcal{V}^h$ as

$$
\begin{aligned}
\mathcal{S}^h = \{\rho \mid \rho(\cdot,t) \in \mathcal{H}^1(\Omega), \rho(\cdot,t)_{|\Omega_e} \in \mathcal{P}_m(\Omega_e),\ t \in [0,T]\ \forall e \text{ and } \rho = \rho^D \text{ on } \Gamma_D\} \\
\mathcal{V}^h = \{\omega \in \mathcal{H}^1(\Omega)\ , \omega_{|\Omega_e} \in \mathcal{P}_m(\Omega_e)\ \forall e \text{ and } \omega = 0 \text{ on } \Gamma_D\},
\end{aligned}
\tag{22}
$$

where $\mathcal{P}_m$ is the finite element interpolating space, for which we use polynomials of order one.

The problem reduces now to find $\rho^h \in \mathcal{S}^h$ for any $t \in [0,T]$ for all $\omega^h \in \mathcal{V}^h$

$$
(\omega^h, \frac{\rho^h}{\Delta t}) + \theta\hat{\mathcal{L}}^h(\rho^h) = (\omega^h, \frac{\rho^n}{\Delta t}) - (1-\theta)\hat{\mathcal{L}}^h(\rho^n) + (\omega^h, s). \tag{23}
$$

The terms associated with the SUPG stabilization are

$$
\begin{aligned}
S^h(\omega^h,\rho^h) = \mathcal{P}(w)\tau\left[\left(\omega^h, \frac{\rho^h}{\Delta t}\right) + \theta\mathcal{L}^h(\rho^h)\right] \\
S^n(\omega^h,\rho^n) = \mathcal{P}(w)\tau\left[\left(\omega^h, \frac{\rho^n}{\Delta t}\right) + (1-\theta)\mathcal{L}^h(\rho^n) + (\omega^h, s)\right]
\end{aligned}
\tag{24}
$$

It is important to note that a second order differential operator appears in $\mathcal{L}^h(\bullet^h)$.

The domain $\Omega$ is discretized in $n_{el}$ elements $\Omega_e$ and we use an isoparametric interpolation with shape functions $N_I$. In our 1D problem, we use element of same size $h = L(t)/n_{el}$. We define $\mathbb{B} = \{I \,|\, I = 1, n_{np}\}$ the full set of $n_{np}$ global node points in the finite element discretization described as $\mathbb{B} = \bigcup_{e=1}^{n_{en}} \mathbb{B}^e$, where $\mathbb{B}^e$ are the set of all element nodes $n_{en}$, $\mathbb{B}^e = \{i \,|\, i = 1, n_{en}\}$. The geometry is then approximated as

$$
\mathbf{x}^h = \sum_{I=1}^{n_{en}} N_I\, \mathbf{x}_I \tag{25}
$$

where the nodal positions are $\mathbf{x}_i$, $i = 1..n_{en}$ and

$$
\omega^h = N_I \in\ \mathcal{V}^h \quad \text{and} \quad \rho^h = \sum_{J=1}^{n_{en}} N_J \rho_J \in\ \mathcal{S}^h \tag{26}
$$

Considering $\mathbb{B}_D$ the set of nodes on the Dirichlet boundary, and noting that our problem do not include Dirichlet B.C, the approximation $\rho^h$ can be expressed as,

$$\rho^h(\mathbf{x}, t) = \sum_{I \in \mathbb{B}} N_I(\mathbf{x}) \rho_I \tag{27}$$

The standard Galerkin solution presents non-physical oscillations when the Peclet number is larger than one [115], that is for convective dominant problems. In order to avoid instabilities, we introduce the SUPG stabilization (see [115] for more details). We introduce a estabilization term, $S(\omega, \Delta\rho)$, in the residual semi-discretized weak form (15)

$$\mathcal{R}(\rho) = \frac{\rho - \rho^n}{\Delta t} + \theta \mathcal{L}(\rho) + (1-\theta)\mathcal{L}(\rho^n) - s, \tag{28}$$

as

$$S(\omega, \rho) = \sum_e \int_{\Omega_e} \mathcal{P}(\omega) \tau \mathcal{R}(\rho) \mathrm{d}\Omega \tag{29}$$

where $\mathcal{P}(\omega) = v\partial_x \omega$. Note that once we perform the finite element discretization to the stablization term, the computation of $S(\omega, \rho)$ is restricted to each element. The stabilization parameter, $\tau$, is (see [115] for more details):

$$\tau = \left( \frac{2}{\Delta t} + \frac{2|v|}{h} + \frac{4D}{h^2} + \sigma \right)^{-1}. \tag{30}$$

Then, the problem reduces to find $\rho^h \in \mathcal{S}^h$ for any $t \in [0, T]$ for all $\omega^h \in \mathcal{V}^h$

$$(\omega^h, \frac{\rho^h}{\Delta t}) + \theta \hat{\mathcal{L}}^h(\rho^h) + S^h(\omega^h, \rho^h) = (\omega^h, \frac{\rho^n}{\Delta t}) - (1-\theta)\hat{\mathcal{L}}^h(\rho^n) + (\omega^h, s) + S^n(\omega^h, \rho^n). \tag{31}$$

Finally, the full discretized transport equation reads

$$\left[ \frac{\mathbf{M} + \mathring{\mathbf{M}}}{\Delta t} \right] \rho + \theta(\boldsymbol{\mathcal{L}} + \mathring{\boldsymbol{\mathcal{L}}})\rho = \left[ \frac{\mathbf{M} + \mathring{\mathbf{M}}}{\Delta t} \right] \rho^n - (1-\theta)[\boldsymbol{\mathcal{L}} + \mathring{\boldsymbol{\mathcal{L}}}]\rho^n + [\mathbf{m} + \mathring{\mathbf{m}}_{\mathbf{s}}] \tag{32}$$

where

$$\boldsymbol{\mathcal{L}} = \mathbf{m_c} + \mathbf{K} + \sigma \mathbf{M} \text{ and } \mathring{\boldsymbol{\mathcal{L}}} = \mathring{\mathbf{m}}_{\mathbf{c}} + \mathring{\mathbf{K}} + \sigma \mathring{\mathbf{M}} \tag{33}$$

and

$$\mathbf{M} = \mathop{\mathsf{A}}_{e=1}^{n_{el}} \int_{\Omega_e} N_i N_j \mathrm{d}\Omega, \quad \mathring{\mathbf{M}} = \mathop{\mathsf{A}}_{e=1}^{n_{el}} \int_{\Omega_e} \tau v N_i' N_j \mathrm{d}\Omega, \quad \mathbf{m} = \mathop{\mathsf{A}}_{e=1}^{n_{el}} \int_{\Omega_e} N_i \mathrm{d}\Omega, \quad \mathring{\mathbf{m}} = \mathop{\mathsf{A}}_{e=1}^{n_{el}} \int_{\Omega_e} \tau v N_i' \mathrm{d}\Omega,$$
$$\mathbf{m_c} = \mathop{\mathsf{A}}_{e=1}^{n_{el}} \int_{\Omega_e} N_i' v N_j \mathrm{d}\Omega, \quad \mathring{\mathbf{m}}_{\mathbf{c}} = \mathop{\mathsf{A}}_{e=1}^{n_{el}} \int_{\Omega_e} \tau v^2 N_i' N_j \mathrm{d}\Omega \quad \text{and} \quad \mathbf{K} = \mathop{\mathsf{A}}_{e=1}^{n_{el}} \int_{\Omega_e} N_i' D N_j' \mathrm{d}\Omega \tag{34}$$

$\mathop{\mathsf{A}}_{e=1}^{n_{el}} \int_{\Omega_e}$ denotes the assembly operator acting on the local element matrix and nodal vectors.

##### A.2 Momentum-balance equation

Let us recall the balance of linear momentum of our problem as

$$\frac{\partial}{\partial x}\left[\mu \frac{\partial v}{\partial x} + \zeta \rho\right] = \eta v \quad in \Omega \tag{35}$$

$$\begin{array}{ccccc} \mu \partial_x v + \zeta \rho & = & \tau(l_f(t)) & \text{on} & \Gamma_N^f \\ v^F & = & 0 & \text{on} & \Gamma_D^c. \end{array} \tag{36}$$

Recall that $\mu$ the viscous coefficient, $\zeta$ is the contractility constant and $\eta$ is the friction parameter. We derive the weak form of the problem by making use of the same trial and test function spaces as described above and integrate in $\Omega$:

$$\int_\Omega w(\mu v_x + \zeta \rho)_x \, dx = \int_\Omega \eta w v \, dx \tag{37}$$

We then apply integration by parts in the left hand side integral and get endequation

$$-\int_\Omega \mu w_x v_x \, dx - \int_\Omega \eta w v \, dx = \int_\Omega \zeta w_x \rho \, dx - \tau \qquad \forall w \in \mathcal{V} \tag{38}$$

We make use again of the Galerkin finite element discretization as for the transport equation so, after performing the same procedure as above, we write the final discretized algebraic system of equations as

$$[-\mu \mathbf{K} - \eta \mathbf{M}] v = \mathbf{f} \tag{39}$$

where

$$K_{ij} = \int_\Omega N_i' N_j' \, dx, \quad M_{ij} = \int_\Omega N_i N_j \, dx \quad \text{and} \quad f_i = \int_\Omega \zeta N_i' \, dx - \tau_i \tag{40}$$

### B Model parameters

The effect of the model parameters (see Section 1.1-2.5) has been analyzed in numerous works and will not be discussed here (see, among others, [103, 106, 90, 101, 69, 91, 102, 126]). We take most of the values used in our simulations from these previous publications (see Table 1 for details).

### C Analysis of the spreading cell

##### C.1 Models and boundary conditions analysis in the spreading cell

To analyze the different modeling options for the actomyosin network described in Sections 2.1-2.2, we run the full transport models, the model with one single specie for the actin and myosin forms and a third case option in which F-actin is assumed constant and only the bound myosin motors are computed. We report minor differences in cell migration velocities predictions (Fig. S11) in all the three cases, with migration velocities of $\approx 3.5 - 4 \mu m/s$. There are also minimal differences in the F-actin and bound myosin densities. Because the simplified transport model with one form of actin and one of myosin, allows us to compute the acto-myosin changes while reducing the computational cost of the full model, we took this approach in all the simulations.

Then, we run the model with one forms of actin (F-actin) and one form of myosin (bound) with non-zero Neumann boundary conditions in the actin mechanics problem (see Section 2.4). The imposition of a stress in both sides of the cell make the actin velocity to increase (Fig. 12).

| | Spreading/mesenchymal | | Ameboid | |
|---|---|---|---|---|
| | Value | Ref. | Value | Ref. |
| $\mu$ [kPa·s] | 10 | [106, 102] | 6 | [134] |
| $\eta$ [kPa·s/$\mu$m] | 0.1 | [106, 102] | 0.01 | [69, 126] |
| $\zeta$ [kPa] / $G_0^*$[kPa] | 0.05 | [106, 90] | 0.1 | [69, 126] |
| $k_p$ / $k_d$ | 0.1/0.1 | [29, 91] | 0.1/0.1 | * |
| $D_F$[$\mu$m/s] | 0.3 | * | 0.3 | * |
| $D_G$[$\mu$m/s] | 0.8 | * | 0.8 | * |
| $D_M$[$\mu$m/s] | 0.4 | * | 0.4 | * |
| $D_m$[$\mu$m/s] | 0.8 | * | 0.8 | * |
| $D_R$[$\mu$m/s] | 0.1 | * | - | |
| $D_S$[$\mu$m/s] | 10 | * | - | |
| $k$ [nN/$\mu$m] | 0.05 | * | 0.5 | * |
| $v_0^p$ [$\mu$m/s] | 0.55 | [20, 113] | - | |
| $L_{pre}(L_0$ [$\mu$m] | 0.3 | * | 0 | * |

Table 1: Material parameters of all computational simulation performed in the main text. All references were used to obtained magnitudes of the corresponding parameters. * indicates that the values were fitted in this work.

This increase makes the densities of actin and myosin to further polarize with respect to the case of zero stresses in the BC. Because the retrograde flow velocity increases, the spreading process is also smaller compared with the zero stresses case in the BC. The stress in the F-actin change from tension to a compressice state at the cell sides and tension at the cell center. This results indicates that, if stresses are applied at the F-actin boundary, the network change from a compressive to a tensional state.

#### C.2 Stiffness of the substrate modifies cell spreading

When a cell is placed on a substrate, it spreads to balance the active contractile and viscous forces with the friction. The spreading area and rate vary when cells are cultured on substrates of varying rigidity [135, 136, 137], because cells form larger AC and adhere more strongly on stiffer substrates, exerting larger traction forces on the substrate [128]. The size of the FAs increases [47] during cell spreading but they also grow when cells are seeded on substrates of varying rigidity [123, 124, 125]. The spreading area also increases as the rigidity of the substrate increase [135, 136, 137].

In our simulations, we assume that the friction with the substrate increases as the stiffness of the substrate increases [123, 124, 125, 128] because the stronger adhesion complexes formed in stiff substrates creates a higher friction of the F-actin network with the substrate. Therefore, an increasing substrate stiffness is proportional to friction. We perform a series of simulations with varying friction $\eta$, which controls the amount of adhesion complexes between the cell and the extracellular space (Fig. 13). Our results show that the spreading radius and the spreading velocity increases with an increasing friction (Fig. 13a,c) and that they increases at similar rates (Fig. 13b,d). When the friction increases as a results of adhesion complex growth, the retrograde flow velocity is constrained and reduced. Therefore, the decrease in the velocity of the retrograde flow results in the net increase of the spreading velocity and the final radius of the cell.

##### C.3 Inhibition of contractile forces fosters cell spreading

Myosin motors exert forces in the actin network and generate a retrograde flow that directly impacts cell motility. It has been shown that depletion of non-muscle myosin-IIA (NMM-IIA), the myosin involved in those contractile forces, increased not only the spreading rate but also the spread area [122]. To analyze whether our model is capable of reproducing these experimental finding and provide a rationalization, we inhibit myosin activity by reducing the contractile force that the motors can exert. Specifically, we vary the parameter $\zeta$ from its baseline value (see Table 1).

Our results are consistent with previous findings (Fig. 14). We show that the spreading radius increases as the contractility decreases (Fig. 14a). We also show a minor increase in the protruding rate that is maintained along time (Fig. 14b). To better understand the reason of these changes, we plot the the retrograde flow velocity as well as the actin and myosin profiles at steady state (Fig. 14c-d). We show that the retrograde flow increases with increased contractility. Because the velocity of the retrograde flow increases, the spreading velocity is reduced. Moreover as the retrograde flow increases and the spreading velocity decreases, the membrane tension also decreases which, in turn, reduces the increase of the retrograde flow and decrease of polymerization velocity. Together, these results are consistent with previous data showing than myosin inhibition increases membrane tension while increasing contractility reduces it [122] and provides a mechanistic rationale.

#### D Supplementary figures

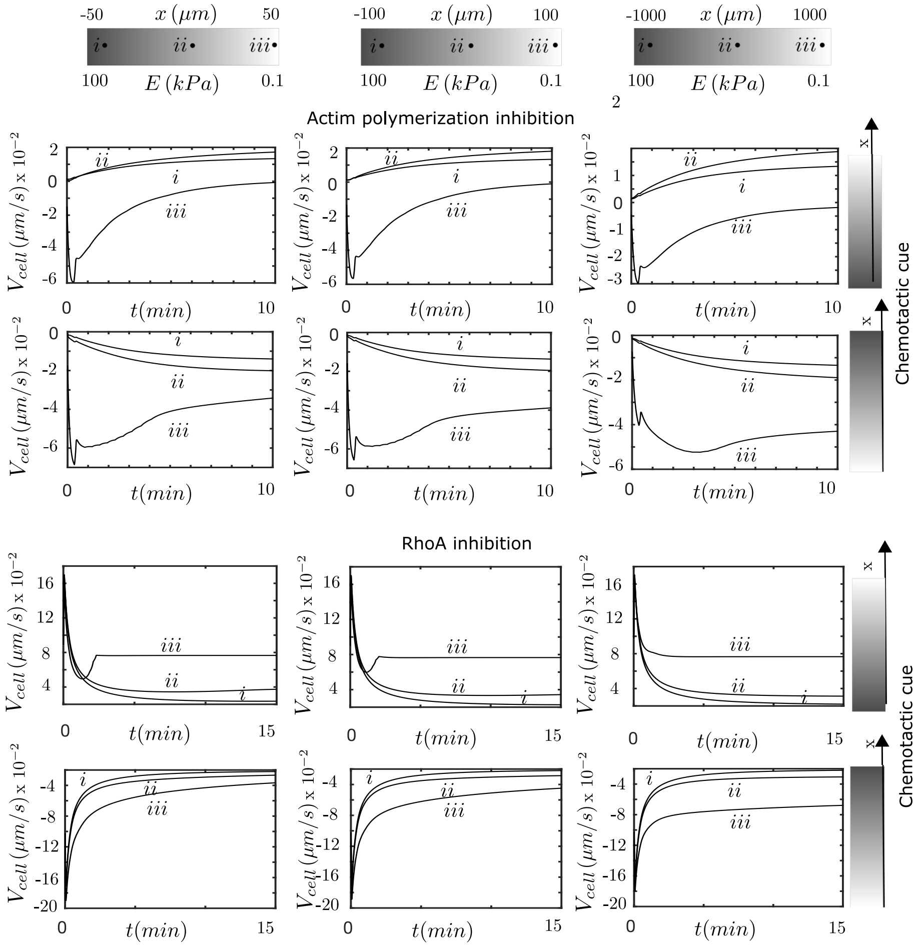


Figure 10: Cell migration velocity under simultaneous durotactic and chemotactic environments. (Top) The polymerization of the cell has been uncoupled from the GTPases signaling and, therefore, it is equal in both cell ends. (Bottom) The myosin activity of the cell has been uncoupled from the GTPases signaling and, therefore, contractibility does not depend on RhoA. Organized by column, samples of 2000, 200 and 100$\mu$m length, from left to right. The substrate stiffness goes from 100kPa, at the right of the sample, to 0.1 kPa, at the left. Points i, ii and ii represent the location where the cells are initially seeded. In first column, samples of 100$\mu m$ in length, i=-30$\mu m$, ii=0$\mu m$, iii=30$\mu m$. In second column, sample of 200$\mu m$ in length, i=-80$\mu m$, ii=0$\mu m$, iii=80$\mu m$. In the third column, samples of 2000$\mu m$ in length, i=-845$\mu m$, ii=0$\mu m$, iii=845$\mu m$. Organized by row, two chemical stimuli in opposing direction. Results in the first row show a chemical cue in the same direction as the durotactic cue. Results in the second row show a chemical cue in the opposite direction to the durotactic cue.

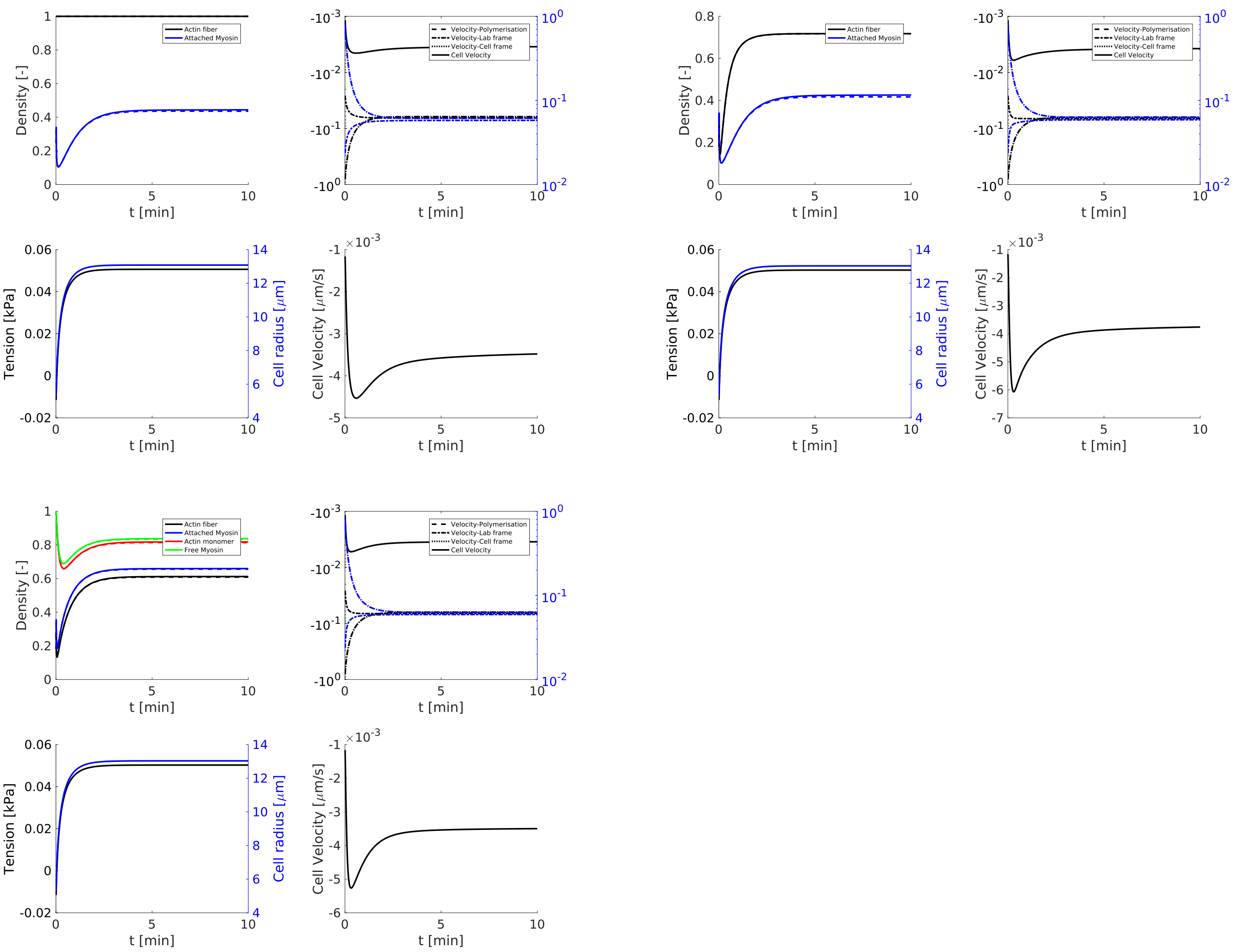


Figure 11: Evolution in time of three transport models of actin and myosin. Top left, model with two forms of actin (F-actin, G-actin) and two forms of myosin (bound and unbound). Top right, model with one forms of actin (F-actin) and one forms of myosin (bound). Bottom, a model with a combined acto-mysoin network. For each model, we show the actin (black) and myosin (blue) densities at the center (dash) and front (solid) of the cell, the polymerization velocity (dot-dash), retrograde velocity at the cell membrane (dash) and total velocity of protrusion (solid), the membrane tension (black) and cell radius (blue) and the migration velocity.

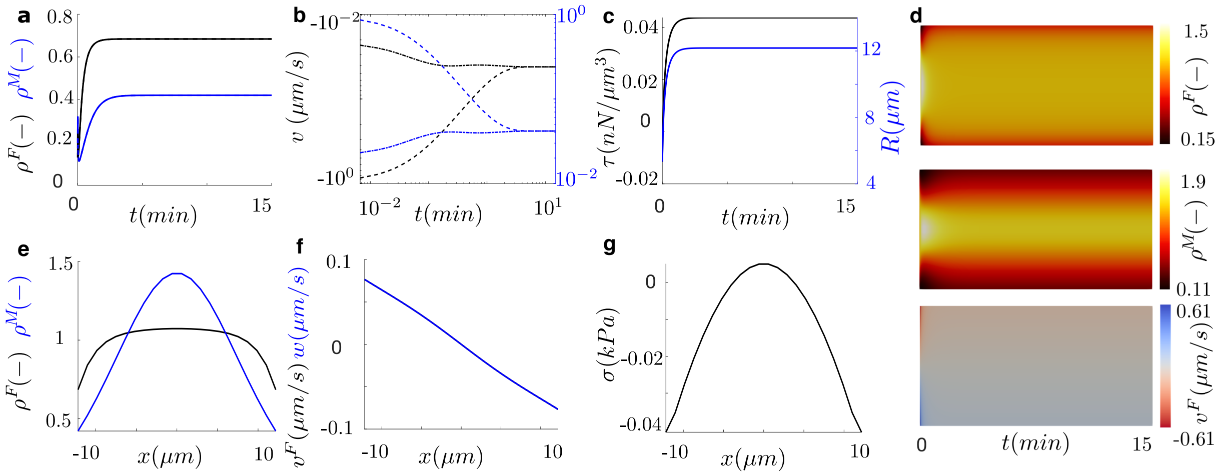


Figure 12: Time and space evolution of cell spreading with cell boundary conditions. (a) Actin (black) and myosin (blue) densities at the left (dash) and right (solid) fronts of the cell. (b) Polymerization velocity (dot-dash), retrograde velocity at the cell membrane (dash) and total velocity of protrusion (solid). (c) Membrane tension (black) and cell radius(blue). (d) Kymographs of the actin density (top), myosin density (center) and retrograde flow velocity (bottom). Actin (black) and myosin (blue) densities (e), retrograde flow (f) and tension of the actin network (g) at steady state along the cell length. Although protrusion velocities are equal on both sides, they slightly decrease as a result of the increase of the membrane tension, which is also the result of the increase of the cell length. In all our simulation, at a larger or smaller degree depending on the case (see Fig. 8), actin and myosin density in the cell polarize due to the friction gradient they experience.

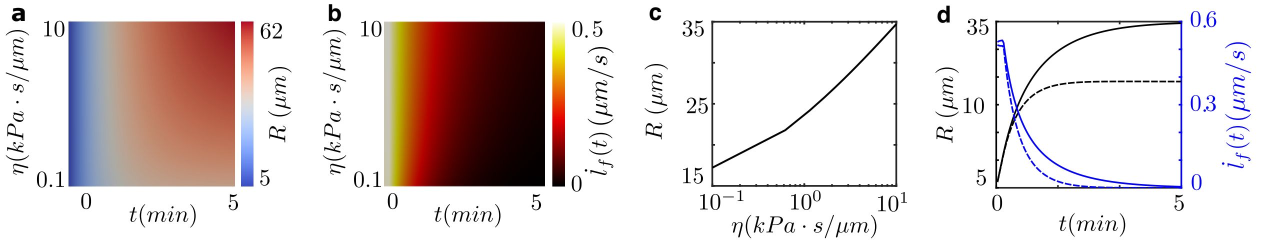


Figure 13: Effect of the substrate rigidity on cell spreading. (a) Evolution of the radius of the cell during 5min as a function of the cell friction with the substrate. (b) Evolution of the cell membrane velocity during 5min. as a function of the cell friction with the substrate. (c) Relation of the final cell radius with the cell friction. (d) Evolution of the radius (black) and cell membrane velocity (blue) in time for $\eta_0 = 1kPa \cdot s/\mu m$ (dash) and $\eta_0 = 10kPa \cdot s/\mu m$.

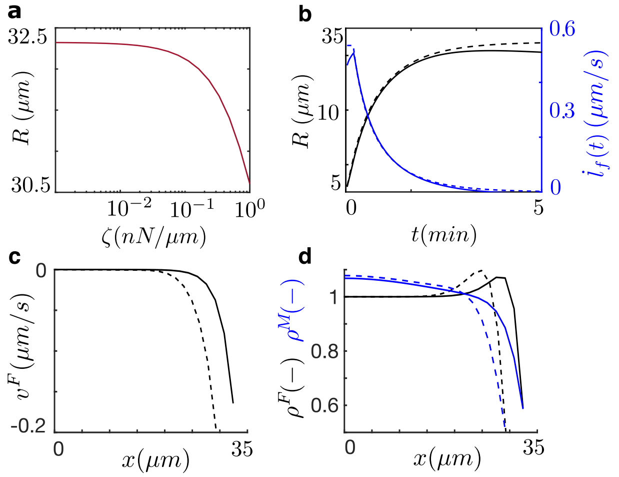

Figure 14: Effect of the substrate rigidity on cell spreading. (a) Evolution of the radius of the cell during 5min as a function of the cell friction with the substrate. (b) Evolution of the cell membrane velocity during 5min. as a function of the cell friction with the substrate. (c) Relation of the final cell radius with the cell friction. (d) Evolution of the radius (black) and cell membrane velocity (blue) in time for $\chi = 0.001 kPa \cdot s/\mu m$ (dash) and $\chi = 1 kPa \cdot s/\mu m$.

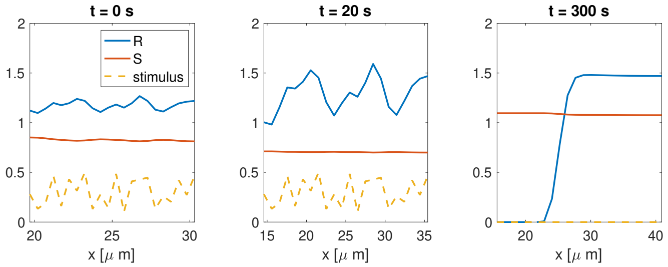


Figure 15: Polarization of active ($\rho^R$) and inactive ($\rho^S$) GTPases due to a random external stimulus (yellow dash) applied at at initial time, t=0. The external stimulus is switched off at t=20s. The steady state polarization of active GTPases is reached after $\approx 8min$.

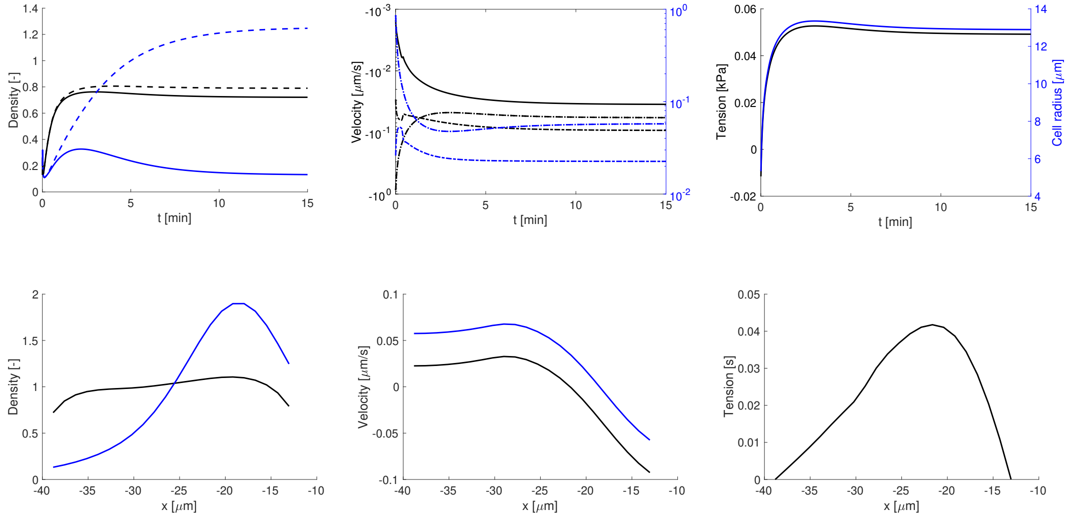


Figure 16: Time and space evolution of the mesenchymal cell migration for inhibition of Rac1. (a) Actin (black) and myosin (blue) densities at the left (dash) and right (solid) fronts of the cell. (b) Polymerization velocity (dot-dash), retrograde velocity at the cell membrane (dash) and total velocity of protrusion (solid). (c) Membrane tension and cell radius. (d) Actin (black) and myosin (blue) densities, (e) retrograde flow and (f) tension of the actin network at steady state along the cell length.

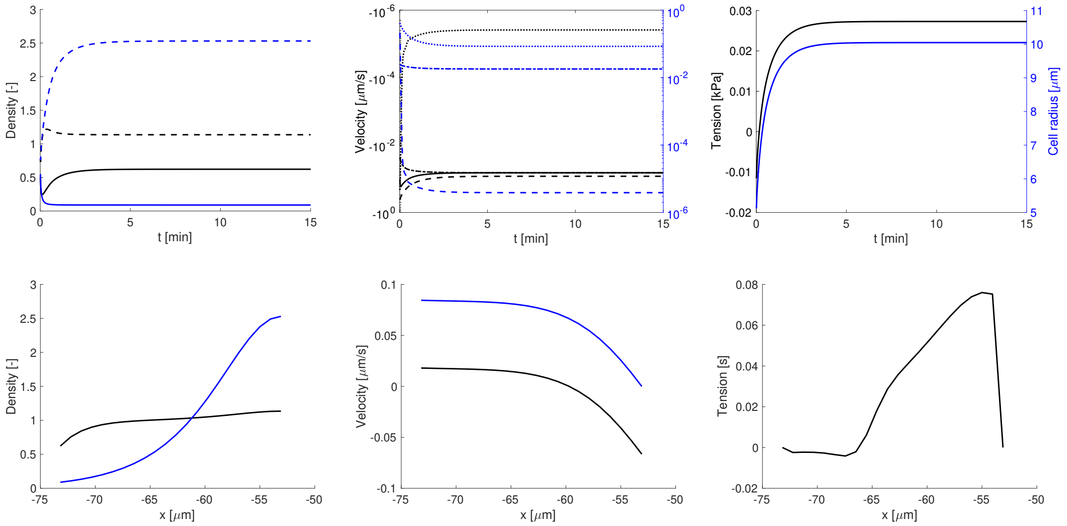


Figure 17: Time and space evolution of the mesenchymal cell migration for inhibition of RhoA. (a) Actin (black) and myosin (blue) densities at the left (dash) and right (solid) fronts of the cell. (b) Polymerization velocity (dot-dash), retrograde velocity at the cell membrane (dash) and total velocity of protrusion (solid). (c) Membrane tension and cell radius. (d) Actin (black) and myosin (blue) densities, (e) retrograde flow and (f) tension of the actin network at steady state along the cell length.

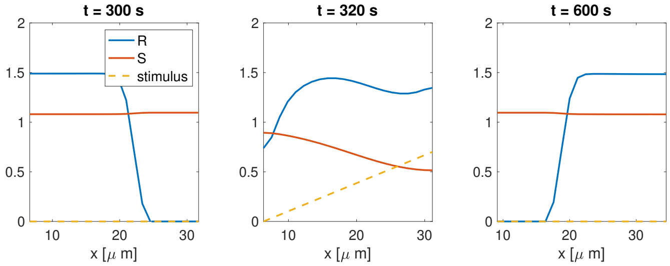


Figure 18: To analyze the sensitivity of cells to new incoming signals, we first induce the polarization of GTPases and compute the solution up to 5s. Then, we add a new gradient stimulus toward the opposite direction of the polarized active GTPases. The GTPases polarizes toward the stimulus direction and a new steady state regime is reached after ≈8min after the new stimulation.

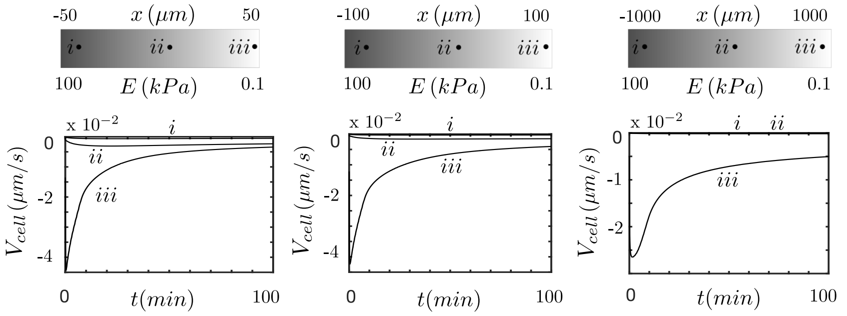


Figure 19: Cell migration velocity at different durotactic environments until steady state, at ≈100min. The substrate stiffness goes from 100kPa, at the left of the sample, to 0.1 kPa, at the right. Points i, ii and ii represent the location where the cells are initially seeded. (a) Sample of 100$\mu m$ in length, i=-30$\mu m$, ii=0$\mu m$, i=30$\mu m$. (b) Sample of 200$\mu m$ in length, i=-80$\mu m$, ii=0$\mu m$, i=80$\mu m$. (c) Sample of 2000$\mu m$ in length, i=-845$\mu m$, ii=0$\mu m$, i=845$\mu m$.

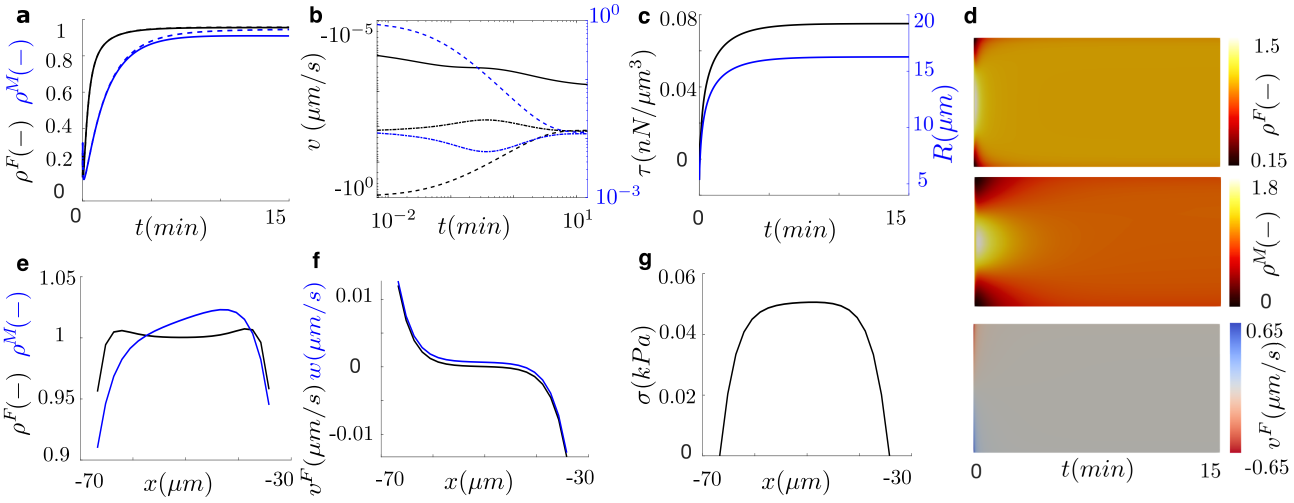


Figure 20: Time and space evolution of cell migration during durotaxis for cell seeded on soft substrates with large stiffness gradient (L=100 $\mu m$). (a) Actin (black) and myosin (blue) densities at the left (dash) and right (solid) fronts of the cell. (b) Polymerization velocity (dot-dash), retrograde velocity at the cell membrane (dash) and total velocity of protrusion (solid). (c) Membrane tension. (d) Kymographs of the actin density (top), myosin density (center) and retrograde flow velocity (bottom). Actin (black) and myosin (blue) densities (e), retrograde flow (f) and tension of the actin network (g) at steady state along the cell length. Although protrusion velocities are equal on both sides, they slightly decrease as a result of the increase of the membrane tension, which is also the result of the increase of the cell length. In all our simulation, at a larger or smaller degree depending on the case (see Fig. 8), actin and myosin density in the cell polarize due to the friction gradient they experience.

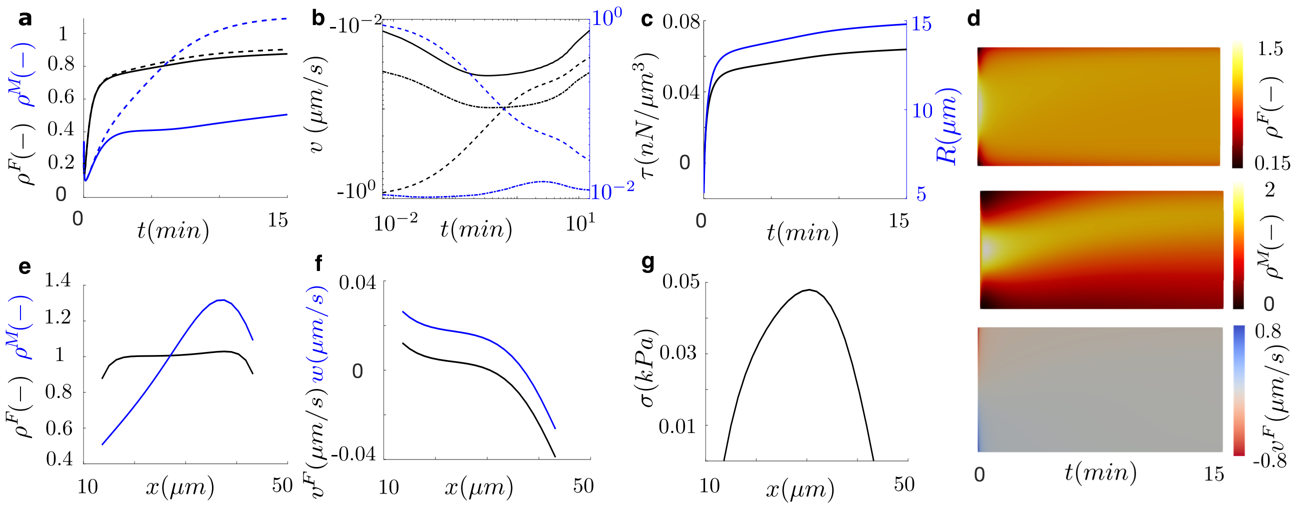


Figure 21: Time and space evolution of cell migration during durotaxis for cell seeded on stiff substrates with large stiffness gradient (L=100 $\mu m$). (a) Actin (black) and myosin (blue) densities at the left (dash) and right (solid) fronts of the cell. (b) Polymerization velocity (dot-dash), retrograde velocity at the cell membrane (dash) and total velocity of protrusion (solid). (c) Membrane tension. (d) Kymographs of the actin density (top), myosin density (center) and retrograde flow velocity (bottom). Actin (black) and myosin (blue) densities (e), retrograde flow (f) and tension of the actin network (g) at steady state along the cell length. Although protrusion velocities are equal on both sides, they slightly decrease as a result of the increase of the membrane tension, which is also the result of the increase of the cell length. In all our simulation, at a larger or smaller degree depending on the case (see Fig. 8), actin and myosin density in the cell polarize due to the friction gradient they experience.

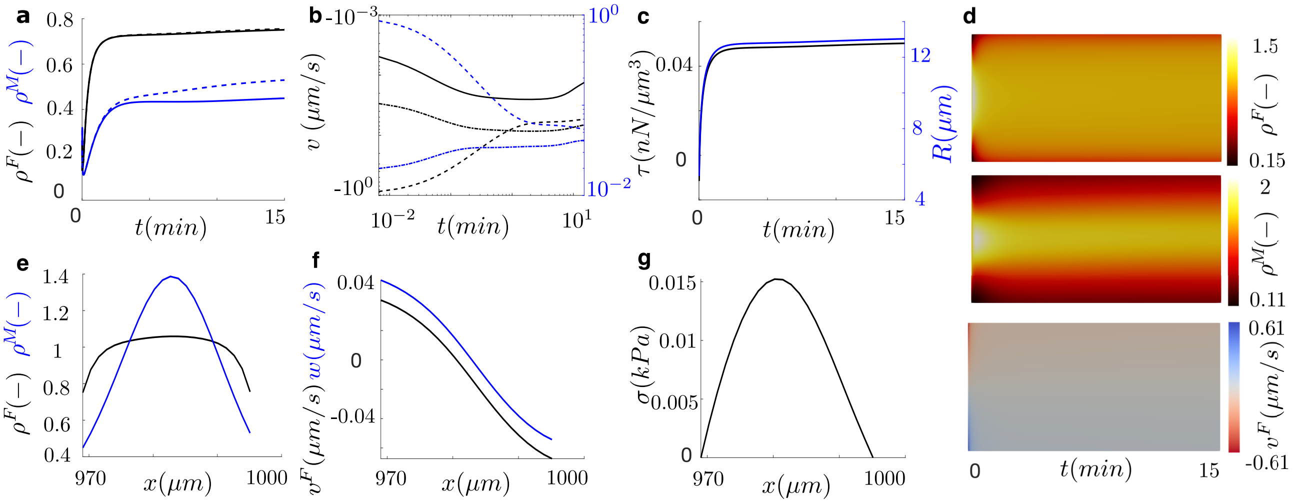


Figure 22: Time and space evolution of cell migration during durotaxis for cell seeded on soft substrates with large stiffness gradient (L=2000 $\mu m$). (a) Actin (black) and myosin (blue) densities at the left (dash) and right (solid) fronts of the cell. (b) Polymerization velocity (dot-dash), retrograde velocity at the cell membrane (dash) and total velocity of protrusion (solid). (c) Membrane tension. (d) Kymographs of the actin density (top), myosin density (center) and retrograde flow velocity (bottom). Actin (black) and myosin (blue) densities (e), retrograde flow (f) and tension of the actin network (g) at steady state along the cell length. Although protrusion velocities are equal on both sides, they slightly decrease as a result of the increase of the membrane tension, which is also the result of the increase of the cell length. In all our simulation, at a larger or smaller degree depending on the case (see Fig. 8), actin and myosin density in the cell polarize due to the friction gradient they experience.

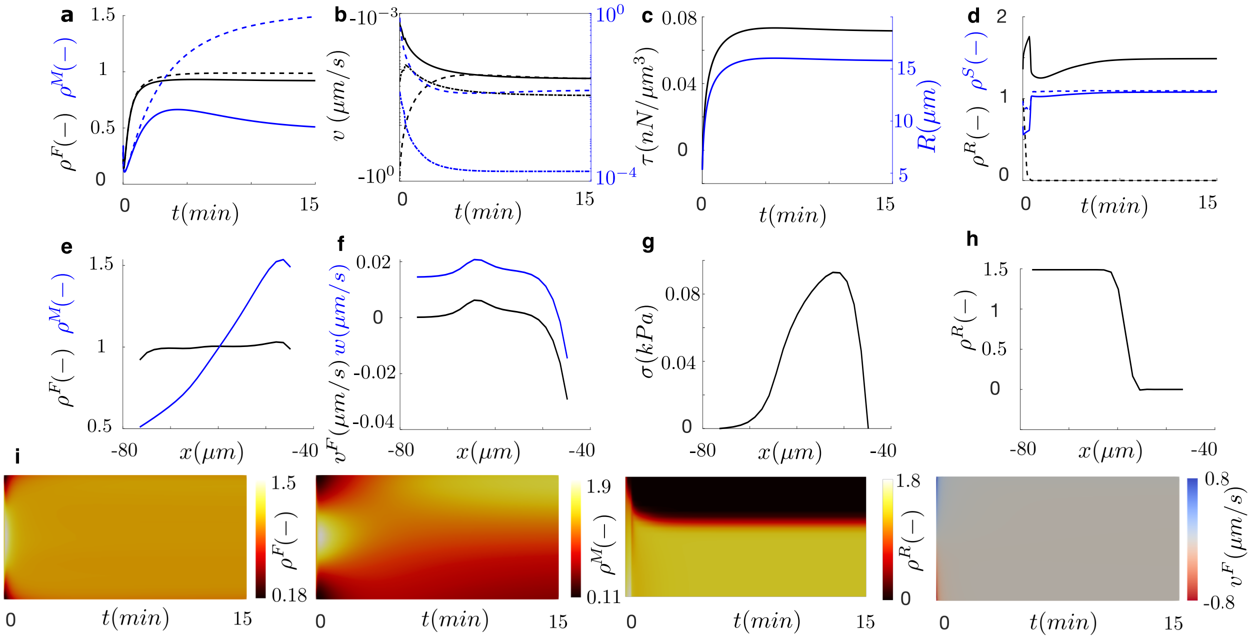


Figure 23: Model results of a cell migrating under cooperative chemical and mechanical stimuli. The cell is placed in a sample of $100\mu m$ in length at the stiffest side of the sample. (a-d) Time evolution of cell spreading. (a) Actin (black) and myosin (blue) densities at the center (dash) and front (solid) of the cell. (b) polymerization velocity (dot-dash), retrograde velocity at the cell membrane (dash) and total velocity of protrusion (solid). (c) Membrane tension. (d) Active and inactive GTPases. (e) Actin (black) and myosin (blue) densities, (f) retrograde flow, (g) tension of the actin network and (h) active and inactive GTPases at steady state along the cell length. (i) Kymographs of the actin density (top), myosin density (center) and retrograde flow velocity (bottom).

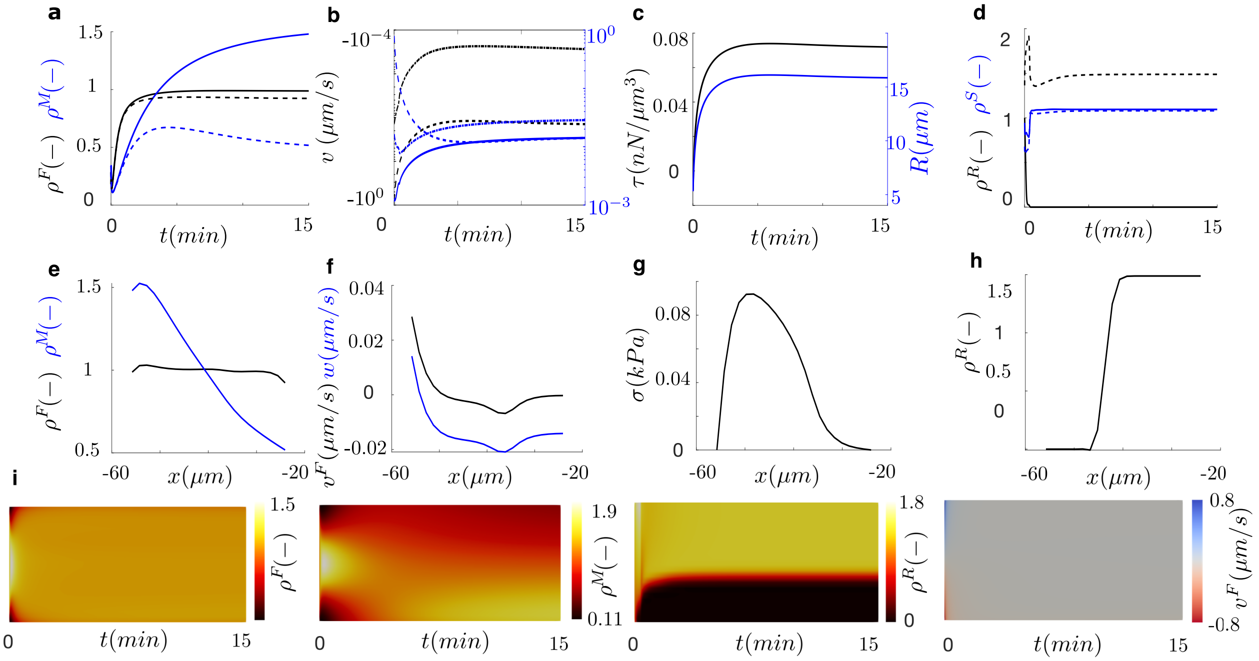


Figure 24: Model results of a cell migrating under cooperative chemical and mechanical stimuli. The cell is placed in a sample of $100\mu m$ in length at the stiffest side of the sample. (a-d) Time evolution of cell spreading. (a) Actin (black) and myosin (blue) densities at the center (dash) and front (solid) of the cell. (b) polymerization velocity (dot-dash), retrograde velocity at the cell membrane (dash) and total velocity of protrusion (solid). (c) Membrane tension. (d) Active and inactive GTPases. (e) Actin (black) and myosin (blue) densities, (f) retrograde flow, (g) tension of the actin network and (h) active and inactive GTPases at steady state along the cell length. (i) Kymographs of the actin density (top), myosin density (center) and retrograde flow velocity (bottom).

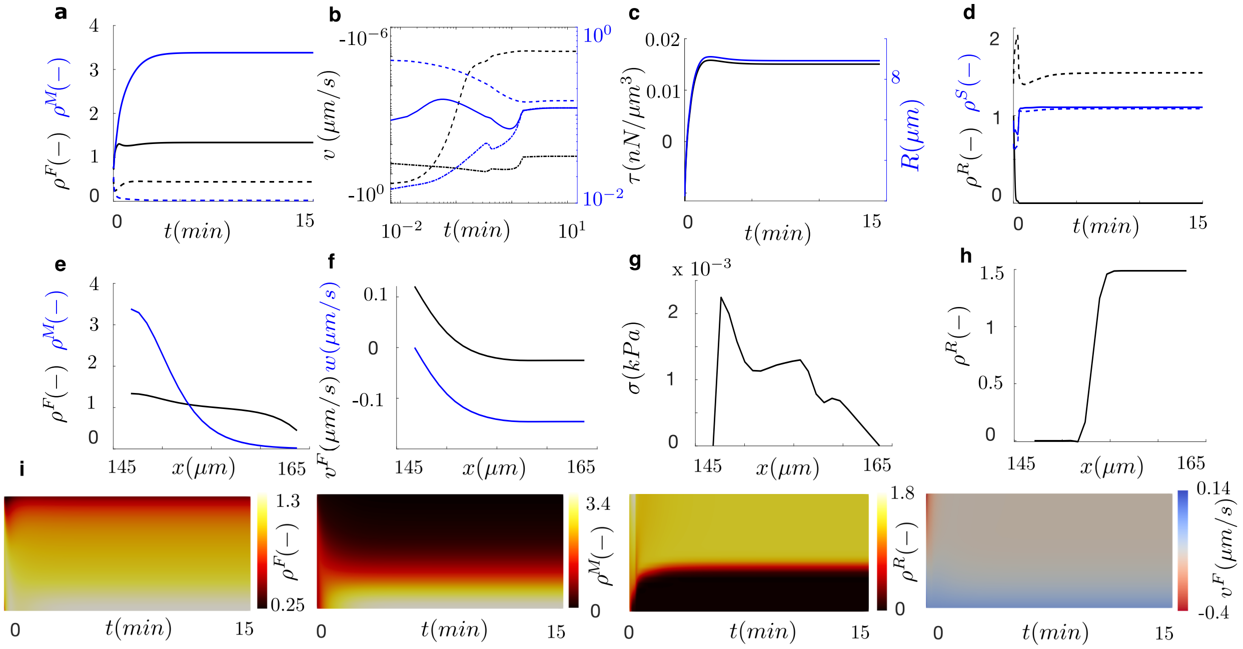


Figure 25: Model results of a cell migrating under cooperative chemical and mechanical stimuli. The cell is placed in a sample of $100\mu m$ in length at the softest side of the sample. (a-d) Time evolution of cell spreading. (a) Actin (black) and myosin (blue) densities at the center (dash) and front (solid) of the cell. (b) polymerization velocity (dot-dash), retrograde velocity at the cell membrane (dash) and total velocity of protrusion (solid). (c) Membrane tension. (d) Active and inactive GTPases. (e) Actin (black) and myosin (blue) densities, (f) retrograde flow, (g) tension of the actin network and (h) active and inactive GTPases at steady state along the cell length. (i) Kymographs of the actin density (top), myosin density (center) and retrograde flow velocity (bottom).

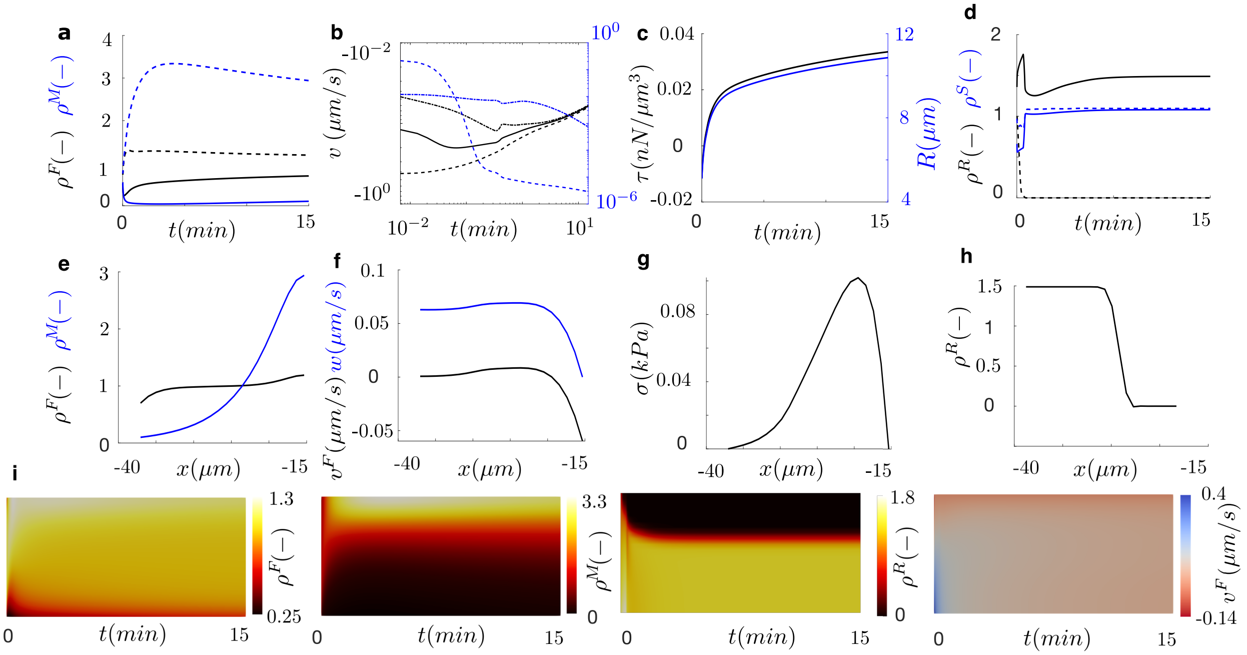


Figure 26: Model results of a cell migrating under cooperative chemical and mechanical stimuli. The cell is placed in a sample of $100\mu m$ in length at the softest side of the sample. (a-d) Time evolution of cell spreading. (a) Actin (black) and myosin (blue) densities at the center (dash) and front (solid) of the cell. (b) polymerization velocity (dot-dash), retrograde velocity at the cell membrane (dash) and total velocity of protrusion (solid). (c) Membrane tension. (d) Active and inactive GTPases. (e) Actin (black) and myosin (blue) densities, (f) retrograde flow, (g) tension of the actin network and (h) active and inactive GTPases at steady state along the cell length. (i) Kymographs of the actin density (top), myosin density (center) and retrograde flow velocity (bottom).

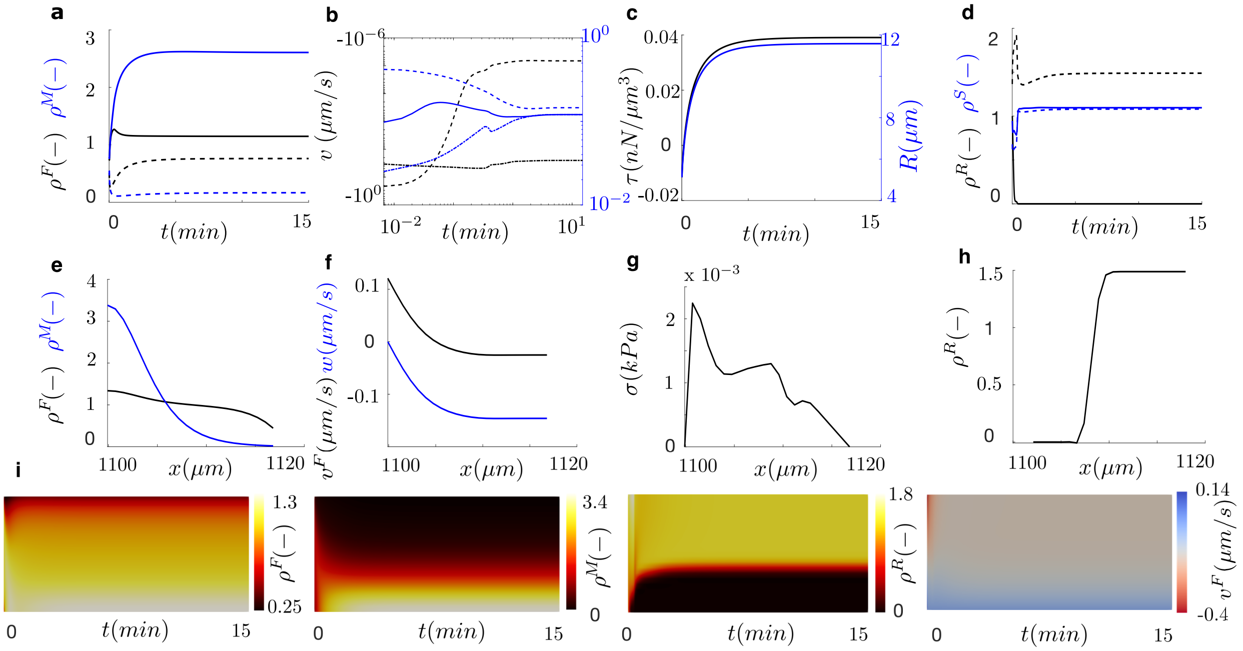


Figure 27: Model results of a cell migrating under cooperative chemical and mechanical stimuli. The cell is placed in a sample of $2000\mu m$ in length at the softest side of the sample. (a-d) Time evolution of cell spreading. (a) Actin (black) and myosin (blue) densities at the center (dash) and front (solid) of the cell. (b) polymerization velocity (dot-dash), retrograde velocity at the cell membrane (dash) and total velocity of protrusion (solid). (c) Membrane tension. (d) Active and inactive GTPases. (e) Actin (black) and myosin (blue) densities, (f) retrograde flow, (g) tension of the actin network and (h) active and inactive GTPases at steady state along the cell length. (i) Kymographs of the actin density (top), myosin density (center) and retrograde flow velocity (bottom).

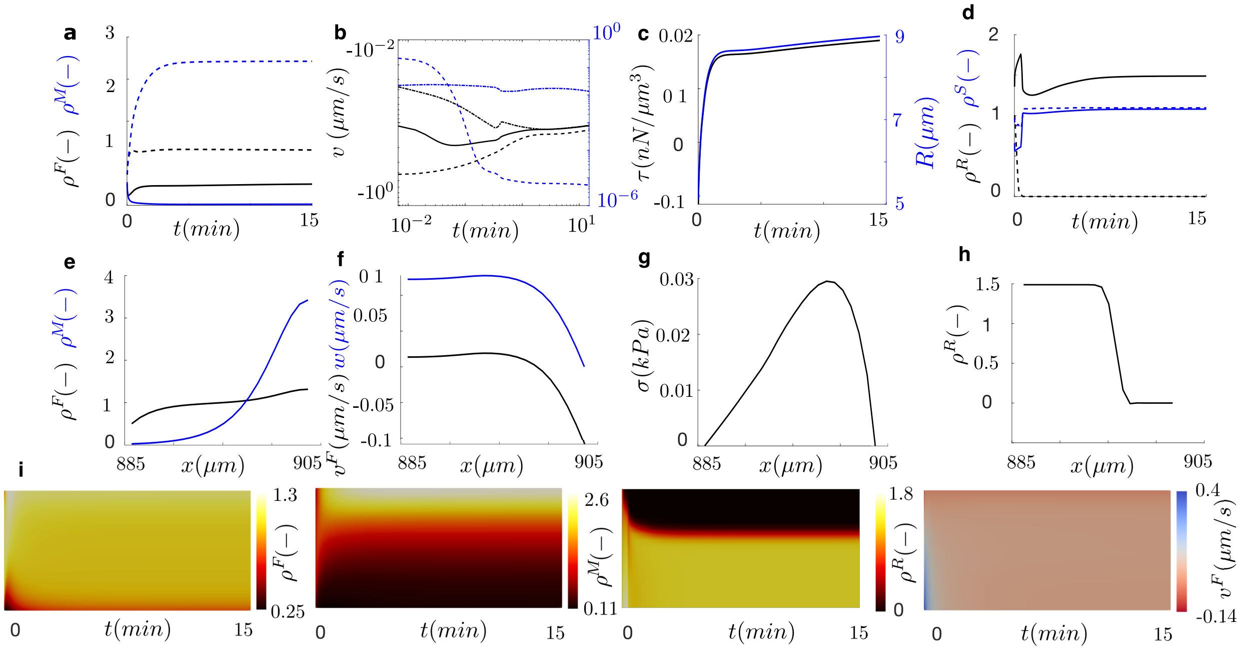

Figure 28: Model results of a cell migrating under cooperative chemical and mechanical stimuli. The cell is placed in a sample of $2000\mu m$ in length at the softest side of the sample. (a-d) Time evolution of cell spreading. (a) Actin (black) and myosin (blue) densities at the center (dash) and front (solid) of the cell. (b) polymerization velocity (dot-dash), retrograde velocity at the cell membrane (dash) and total velocity of protrusion (solid). (c) Membrane tension. (d) Active and inactive GTPases. (e) Actin (black) and myosin (blue) densities, (f) retrograde flow, (g) tension of the actin network and (h) active and inactive GTPases at steady state along the cell length. (i) Kymographs of the actin density (top), myosin density (center) and retrograde flow velocity (bottom).

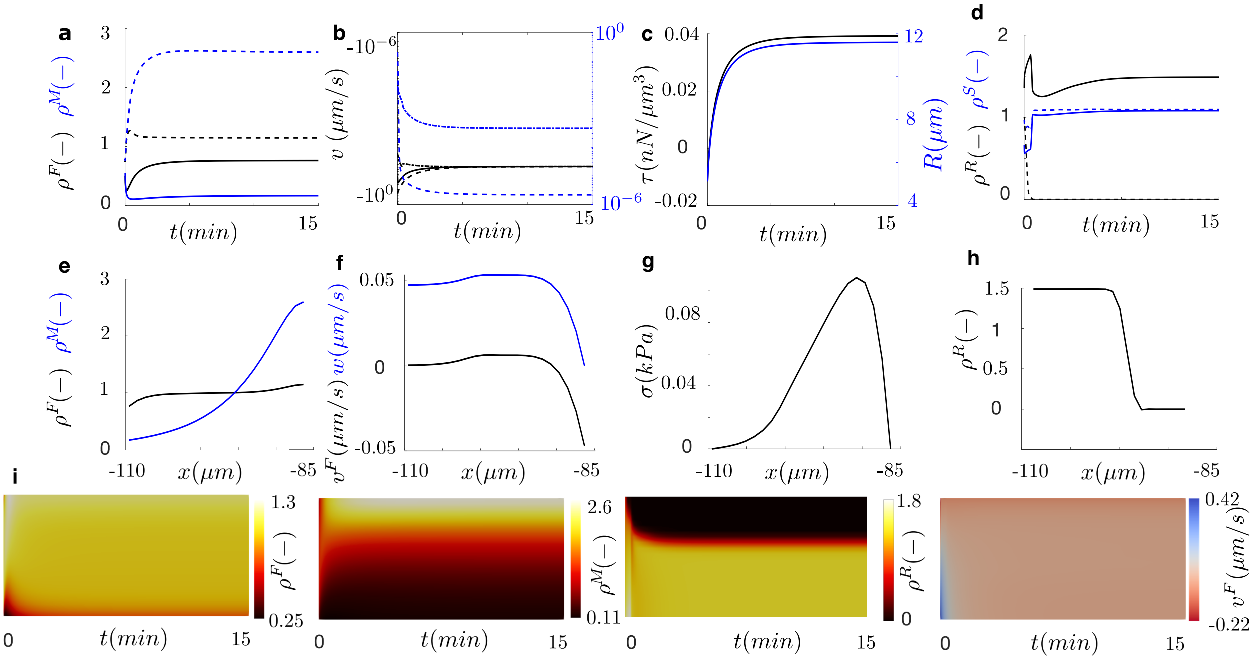


Figure 29: Model results of a cell migrating under cooperative chemical and mechanical stimuli. The cell is placed in a sample of $100\mu m$ in length at the softest side of the sample. The actin polymerization at the cell front has been inhibited. (a-d) Time evolution of cell spreading. (a) Actin (black) and myosin (blue) densities at the center (dash) and front (solid) of the cell. (b) polymerization velocity (dot-dash), retrograde velocity at the cell membrane (dash) and total velocity of protrusion (solid). (c) Membrane tension. (d) Active and inactive GTPases. (e) Actin (black) and myosin (blue) densities, (f) retrograde flow, (g) tension of the actin network and (h) active and inactive GTPases at steady state along the cell length. (i) Kymographs of the actin density (top), myosin density (center) and retrograde flow velocity (bottom).

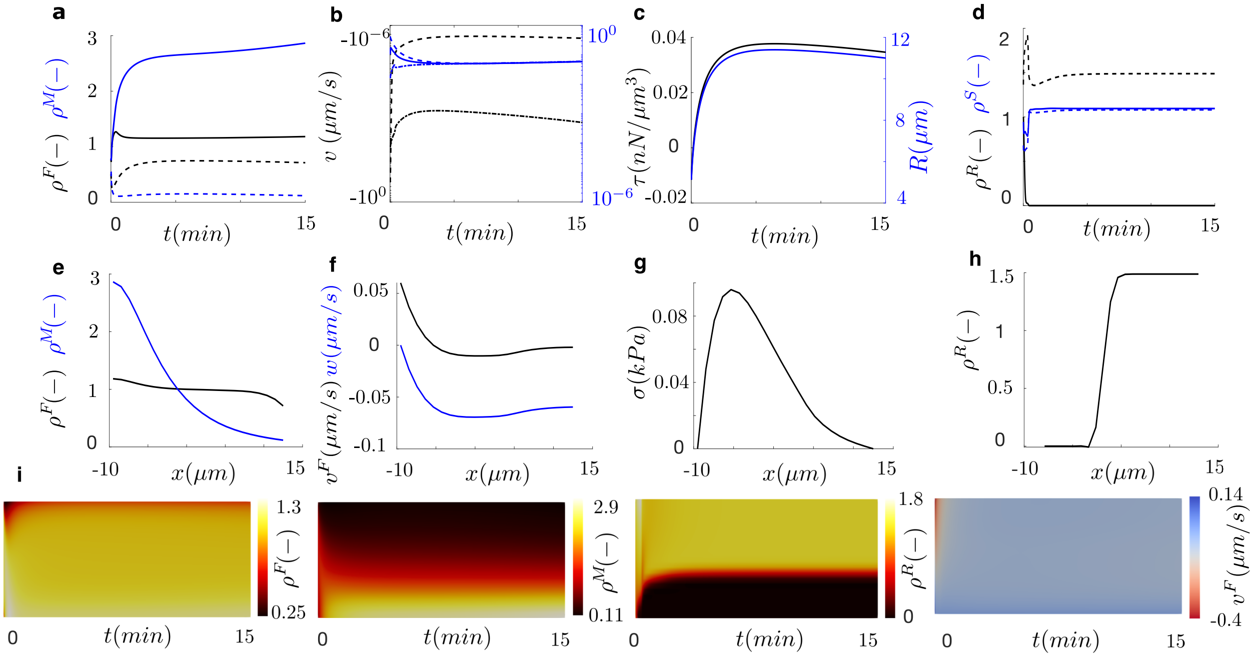

Figure 30: Model results of a cell migrating under competing chemical and mechanical stimuli. The cell is placed in a sample of $100\mu m$ in length at the softest side of the sample. The actin polymerization at the cell front has been inhibited. (a-d) Time evolution of cell spreading. (a) Actin (black) and myosin (blue) densities at the center (dash) and front (solid) of the cell. (b) polymerization velocity (dot-dash), retrograde velocity at the cell membrane (dash) and total velocity of protrusion (solid). (c) Membrane tension. (d) Active and inactive GTPases. (e) Actin (black) and myosin (blue) densities, (f) retrograde flow, (g) tension of the actin network and (h) active and inactive GTPases at steady state along the cell length. (i) Kymographs of the actin density (top), myosin density (center) and retrograde flow velocity (bottom).
